# Enhancing Biomedical Optical Volumetric Imaging via Self-Supervised Orthogonal Learning

**DOI:** 10.1101/2025.05.16.654259

**Authors:** Yuanjie Gu, Yiqun Wang, Ang Xuan, Jianping Wang, Linyi Wang, Lei Zhang, Xiaoran Li, Yao Wu, Jun Zhang, Zhi Lu, Biqin Dong

## Abstract

Optical volumetric imaging grapples with inherent noise problems arising from photon budget constraints, light scattering, and space-bandwidth product bottlenecks, all of which degrade structural fidelity and become even more pronounced than in planar imaging. While deep learning presents potential for denoising, supervised methods are hindered by the impracticality of paired dataset, and existing self-supervised approaches fail to fully exploit the intrinsic volumetric structural redundancy inherent to optical imaging. Here, we present a self-supervised Volumetric biomedicAL Imaging Denoiser (VALID) that leverages intrinsic three-dimensional spatial coherence for highly efficient volumetric denoising through a self-supervised orthogonal learning framework. VALID demonstrates robust denoising performance across diverse imaging modalities, including two- and three-photon microscopy, light-field microscopy, and optical coherence tomography, substantially enhancing structural fidelity in deep-tissue, multimodal, and dynamic imaging scenarios. By combining computational efficiency with zero-shot adaptability, VALID establishes a transformative approach to volumetric image enhancement with structure-aware precision.

## Introduction

Optical volumetric imaging has become indispensable in biomedical research and clinical diagnostics, enabling high-resolution three-dimensional visualization of cellular and tissue architectures in living systems^1–9^. However, these techniques are constrained by fundamental physical limitations: phototoxicity thresholds impose stringent photon budget restrictions, while light scattering during image acquisition degrades signal integrity, introducing noise and information loss in volumetric datasets^10–12^.Furthermore, volumetric imaging systems confront amplified technical challenges compared to conventional two-dimensional imaging, including the space-bandwidth product (SBP) bottleneck^13^, intricate scanning protocol requirement^14^, and artifact generation inherent to volumetric acquisition methodologies^15^. These compounding factors not only collectively undermine biologically relevant structural details and compromise diagnostic reliability but also propagate systematic errors through downstream analytical pipelines that are critical for quantitative biomedical studies^16–18^. This critical landscape underscores the urgent need for advanced denoising algorithms specifically optimized for enhancing volumetric imaging quality.

Recent advances in deep learning-based denoising approaches have demonstrated remarkable capabilities, yet notable limitations remain. Conventional methods relying on handcrafted priors struggle to adequately represent the complex heterogeneity of biological systems, whereas supervised learning approaches necessitate precisely registered noisy-clean image pairs^19–21^, a requirement that proves particularly demanding for volumetric imaging, due to the exorbitant costs of longitudinal imaging protocols and the technical complexity of sample preparation. This fundamental limitation highlights the critical need for developing self-supervised denoising frameworks that can overcome these experimental constraints^22–31^. Fortunately, volumetric imaging data exhibit inherent spatial coherence, where three-dimensional frames maintain consistent structural patterns across sequential acquisitions, with variations primarily arising from stochastic noise. This intrinsic data redundancy provides a foundation for a self-supervised learning framework to discriminatively recover authentic signals by modelling noise distributions in the feature space. The integration of such self-supervised denoising modules into conventional optical volumetric imaging systems represents a pivotal advancement toward intelligent imaging platforms capable of preserving structural fidelity, enhancing signal recovery, and overcoming current experimental limitations.

Several innovative approaches have emerged to address the above persistent challenges through various strategies including spatial redundancy exploitation in single frame^24^, time-lapse blind-spot interpolation by neighbor frames^26,28^, temporal redundancy exploitation using odd-even sampled multi-frame ^29,30^, and adjustable frame-multiplexed sampling to balance the spatial and temporal (Z-axis in volumetric imaging) specific inherent priors^31^. Despite significant progress in self-supervised denoising approaches that exploit two-dimensional spatial and temporal redundancies, current methods remain fundamentally limited in their ability to harness the full three-dimensional information redundancy inherent in volumetric imaging data, a critical capability for accurate visualization of dynamic biological processes in vivo. Conceptually, existing methodologies can be classified into three primary categories (Fig. 1a) based on their perceptual domains, including single-frame perception field, blind-frame perception field, and multi-frame perception field. While each approach serves specific application needs (single-image denoising, ultra-fast time-series denoising, and precise signal extraction, etc.), they collectively fail to fully exploit the integrated three-dimensional structural information where all spatial dimensions (X, Y, Z) jointly characterize the complete sample representation. Therefore, it remains unexplored to unify these perspectives through a volume perception field that inherently captures three-dimensional structural redundancy, enabling a self-supervised framework tailored for volumetric imaging.

**Fig. 1.**
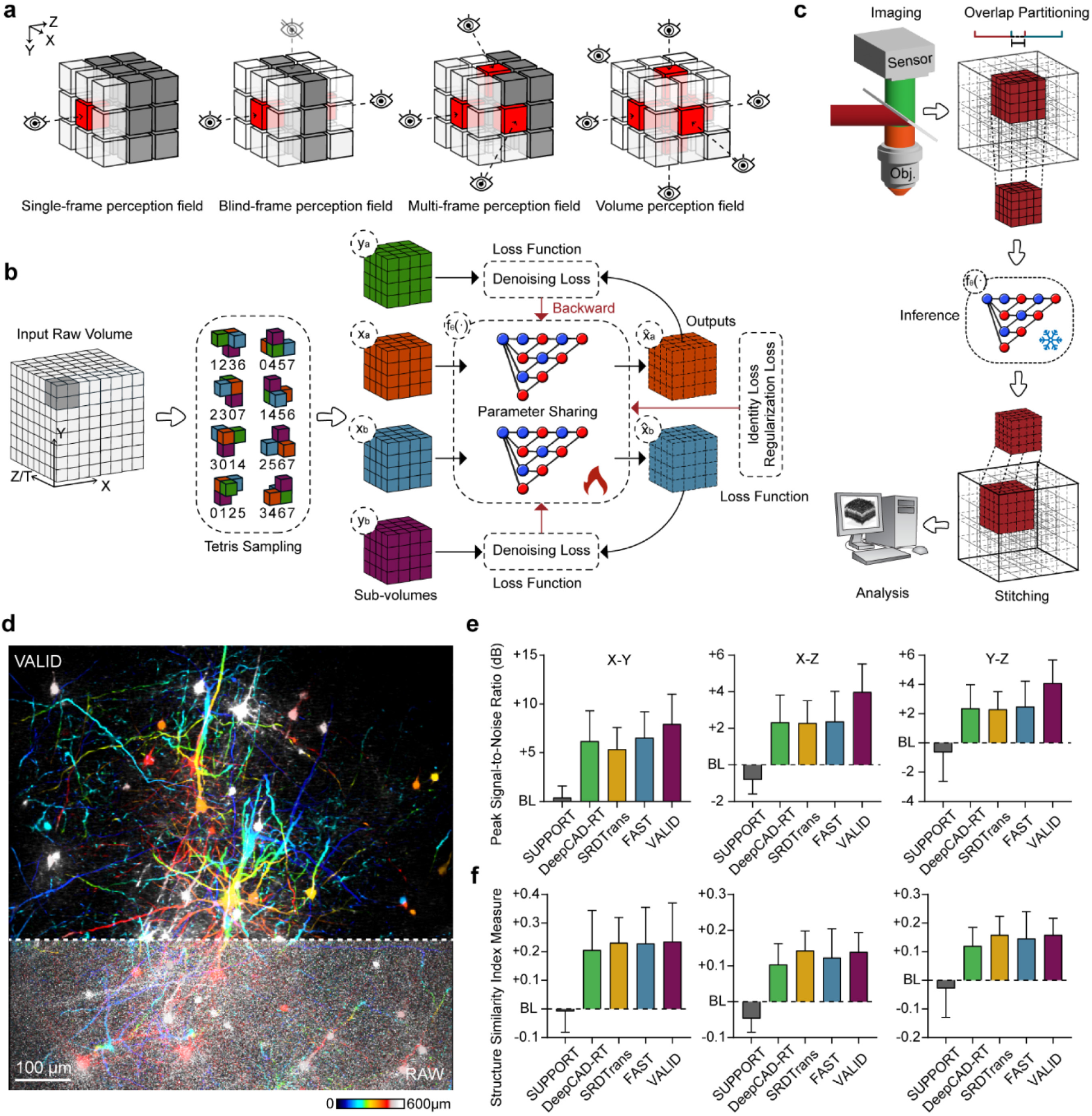
Overview of VALID framework and its application in two-photon volumetric structural imaging. **a,** The conceptual framework of self-supervised denoising strategies comprises four distinct perceptual domains: Single-frame perception leverages intra-frame spatial redundancy through adjacent spatial information within individual frames; Blind-spot perception employs localized redundancy via center-pixel prediction while masking surrounding pixels in single frames; Multi-frame perception exploits inter-frame spatial redundancy by aggregating adjacent spatial information across sequential frames; Volumetric perception integrates comprehensive redundancy through holistic analysis of entire 3D volumes. **b,** VALID’s self-supervised training strategy. Raw low SNR volumetric data (XYZ) or time-laspe data (XYT) undergo Tetris-based initial sampling, dividing the volume into sub-volumes x_a_, x_b_, y_a_, y_b_. Each block within the volume is sampled using one of eight predefined patterns, producing paired sub-volumes where x_a_, y_a_ and x_b_, y_b_ serve as input-target pairs for network optimization. The CRN processes the inputs with shared weights, and optimization is achieved through denoising, identity, and regularization loss functions. **c,** VALID’s inference pipeline. Input volumetric data are partitioned into overlapping 3D patches using a sliding window strategy. Each patch is processed independently by the trained CRN to produce enhanced sub-volumes. The sub-volumes are then reassembled using weighted averaging at overlapping regions to restore the original volume. **d,** Two-photon volumetric structural imaging results. Depth-encoded projection of two-photon imaging data (depth range: 0–600 µm) show raw data (top-left), FAST results (bottom-right), and VALID-enhanced results (top-right). VALID demonstrates improved structural detail and reduced background noise, particularly in deeper regions (400–600 µm). Scale bar, 100 μm. **e**-**f,** Quantitative benchmark of denoising performance with state-of-the-art methods, including SUPPORT, DeepCAD-RT, SRDTrans and FAST. Layer-wise Peak Signal-to-Noise Ratio (PSNR) and Structural Similarity Index Measure (SSIM) values are plotted for comparisons across imaging depths for different planes (X-Y, X-Z and Y-Z). The raw data scores are indicated as the baseline (BL). VALID achieves the best PSNR performance (+8.0 dB, +4.0 dB, and +4.1 dB), improving 23.1%, 68.1%, and 63.9% compared to FAST, while maintaining the competitive SSIM scores.

Here, we propose a structure-friendly volumetric biomedical imaging denoiser, named VALID, to further push the ceilings of biomedical imaging. A deep learning framework designed to overcome current limitations in biomedical volumetric imaging. VALID proposes a novel self-supervised volumetric orthogonal learning strategy, employing Tetris sampling to utilize the cyclic consistency between orthogonal acquisition planes (X-Y, X-Z, Y-Z) to reinforce structural authenticity, and a fine-designed Cross-scale Recursive Network (CRN), while regularized training on noise Hessian matrix prevents signal hallucination. Validation across diverse volumetric biomedical optical imaging techniques (two-, three-photon microscopy, light-field microscopy, and optical coherence tomography) demonstrates significant improvements over conventional methods, particularly in challenging imaging scenarios characterized by extremely low Signal-to-Noise Ratio (SNR) and complex mixed noise distributions. This advancement establishes a new paradigm for intelligent volumetric imaging enhancement, offering a transformative step in biomedical imaging and thereby facilitate more accurate researches in life sciences and clinical diagnostics.

## Results

### Principle of VALID

To address the volume perception gap in denoising for VALID, we propose self-supervised orthogonal learning, a novel strategy that systematically incorporates 3D structural priors (see details in Methods). Additionally, we introduce dimension-aware orthogonal sampling alongside a carefully designed Cross-scale Recursive Network (CRN) to enhance volumetric feature extraction (see Supplementary Fig. 1 and Methods). VALID’s self-supervised orthogonal learning framework operates through two distinct operational stages: training and inference. The training phase follows a structured pipeline (Fig. 1b). Initial Tetris sampling is performed on raw volumetric inputs (either XYZ volumetric spatial or XYT time-lapse spatiotemporal data) to generate paired training sub-volumes {*X*_*a*_, *X*_*b*_} and {*Y*_*a*_, *Y*_*b*_}. These sub-volume pairs are then processed through our specifically designed CRN architecture. The network receives {*X*_*a*_, *X*_*b*_} as input features while {*Y*_*a*_, *Y*_*b*_ } are designated as target outputs for loss computation. During forward propagation, CRN generates predicted sub-volumes 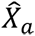 and 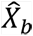, which are subsequently compared with their corresponding target sub-volumes through our multi-objective loss function (consist of denoising, identity, and regularization; see details in Methods). To further enhance the constraints on unconventional noises, such as speckle noise in OCT, we have designed a three-dimensional low-frequency Hessian matrix constraint loss for the orthogonality validation of low-frequency components as a regularization term. Benefiting to the above efforts, the statistically redundant orthogonal features are exploited to drive parameter optimization. It is worth noting that the two inputs *X*_*a*_ and *X*_*b*_ are fed forward into the network that has shared in terms of weights. They do not use two separate sets of parameters for optimization. Thus, during inference, we only need to perform one set of parameters.

Beyond the self-supervised orthogonal learning framework, the carefully crafted CRN architecture stands as another central contribution. While U-Net and its variants have become backbone architectures in the field^32–34^, we observe a fundamental discrepancy: the autoencoder architecture of U-Net was originally designed for high-dimensional semantic feature perception^35^, whereas denoising constitutes a low-dimensional visual task. We hold the view that the effectiveness of U-Net in previous denoising applications does not primarily originate from its multi-level encoder-decoder structure with skip connections, as commonly presumed^31^. Instead, we identify that the critical factor lies in the intrinsic low-pass filtering properties induced by the encoder’s down-sampling operations. Therefore, we re-examined the design of the backbone architecture and proposed CRN, a multi-scale recursive structure restoration network that employs direct latent low-pass filtering instead of encoding (see architecture in Methods and Supplementary Fig. 1). This architecture empowers VALID to achieve better denoising performance and structure maintaining simultaneously significantly fewer parameters. Systematic simulation experiments demonstrate that the combination of CRN and orthogonal learning enables VALID to have a broader generalization ability, especially for out-of-domain data (Supplementary Fig. 2 and Fig. 3).

During inference, the trained CRN with frozen parameters processes unseen volumetric data using a streamlined volumetric patch-based workflow. First, the input volume is partitioned into overlapping 3D patches through a sliding window strategy, mirroring the spatial or temporal sampling principles used in training while ensuring continuity at patch boundaries by overlap. Each patch is independently fed into the optimized CRN network for denoising and feature restoration. The model predicts enhanced sub-volumes, which are then systematically recombined into a complete volume using weighted averaging at overlapping regions, effectively suppressing boundary artifacts, and ensuring spatial consistency.

### Enhancing two-photon volumetric structural imaging

In two-photon microscopy systems^36^, diverse noise sources (particularly shot noise, dark current noise, and readout noise) collectively degrade imaging fidelity despite the modality’s unparalleled deep-tissue penetration capacity. The extended photon propagation path through scattering biological media induces progressive signal attenuation, exacerbating SNR deterioration at increasing imaging depths. While elevated excitation power could theoretically enhance SNR, this strategy requires rigorous optimization to reconcile signal detection efficacy with specimen viability. Excessive laser irradiation induces photochemical perturbations through fluorophore photobleaching and cytotoxic effects, mandating precise intensity calibration to preserve both optical signal integrity and biological sample functionality.

To evaluate the denoising performance of our VALID on two-photon volumetric structural imaging with ground truth, which obtained by averaging intensity of 60 frames. we implemented the algorithm on our two-photon microscope that enabled intravital neuron imaging to depths of ∼600 µm in mouse cortex (*Thy1*-GFP-M). The system was imaged by Resonant-Galvo (X-Y) scanning configuration with a frame rate of 30 Hz. Fig. 1d displays depth pseudo-color encoding projections of 0 to 600 μm depth comparing the result of our proposed VALID (top) with raw data (down). The side-by-side visualization highlights enhanced structural clarity and reduced background noise in our approach, particularly in deeper tissue regions (400-600 μm, orange to white). Fig. 1e quantifies the layer-wise Peak Signal-to-Noise Ratio (PSNR) and Structural Similarity Index Measure (SSIM) across all depths for both comparisons. Our method demonstrates consistently higher PSNR and SSIM values (average scores: PSNR of 39.4 ± 6.26 dB and SSIM of 0.87 ± 0.14), with the most significant gains (***, P<0.001), outperforming raw data by ∼24.7% PSNR and ∼42.6% SSIM scores. The quantitative and qualitative assessment focuses on the VALID’s capability to preserve micron-scale structural details while suppressing out-of-focus background signals. These results demonstrate that VALID can specifically address the unique noise characteristics in two-photon fluorescence microscopy, especially for deep tissue penetration-induced non-uniform Poisson-Gaussian mixed noise patterns.

### Empowering multimodal three-photon volumetric structural imaging

Three-photon multimodal structural imaging is a cutting-edge intravital volumetric imaging technique that enables imaging with greater depth (achieving hippocampal region) than two-photon imaging^37^. As the imaging depth increases, it also faces extreme challenges in achieving sufficient SNR and contrast due to the inherent limitations of nonlinear excitation processes. In deep hippocampal imaging, the amount of available photon is drastically reduced by intensified tissue scattering, and result in degraded signal quality. Even with ideal detectors and lasers, the combination of these factors, depth-dependent scattering, nonlinear signal scaling, and strict energy constraints to avoid tissue damage, creates an unavoidable noise-dominated regime, making precise visualization of neural structures at these depths exceptionally demanding. The extreme low-photon environment in the hippocampus amplifies these challenges^38,39^, necessitating advanced noise suppression and signal optimization strategies to resolve subtle structural and functional details.

To evaluate VALID’s denoising performance in advanced three-photon intravital multimodal imaging of the mouse brain, particularly targeting the hippocampal region from 800 to 1100 μm depths, we implemented VALID on our three-photon microscope (Fig. 2a) featuring a Galvo-Galvo (X-Y) scanning configuration. This system achieved multimodal imaging through synchronized acquisition of vessel (Texas Red), neuron structure (jGCaMP7s), and label-free third harmonic generation (THG) imaging. The pixel SNR values are calculated with 9 slices at the same position (see Methods). Figs. 2b-d demonstrate significant improvements across different comparisons (RAW, FAST, and VALID) in three distinct modalities. For blood vessels, average SNR increased progressively from 1.47 ± 0.22 dB (RAW) to 10.37 ± 0.30 dB (FAST) and further to 14.97 ± 0.68 dB (VALID), enhancing about 10.2-fold SNR against RAW. Similar enhancement patterns were observed in neuronal imaging, with SNR values rising from 1.30 ± 0.32 dB (RAW) through 11.33 ± 0.47 dB (FAST) to 15.49 ± 0.75 dB (VALID), enhancing about 11.9-fold SNR against RAW. THG signals showed the highest baseline performance in RAW mode (2.03 ± 0.18 dB), which was subsequently improved to 11.65 ± 0.32 dB by FAST and ultimately 15.96 ± 0.84 dB by VALID, enhancing about 7.9-fold SNR against RAW. Notably, the VALID framework consistently achieved the highest SNR values across all measured categories, with neuronal signals demonstrating the best overall SNR performance followed by neuronal vasculature and label-free THG’s results. These quantitative results highlight the substantial SNR gains achieved through VALID performing in three-photon microscopy applications.

**Fig. 2.**
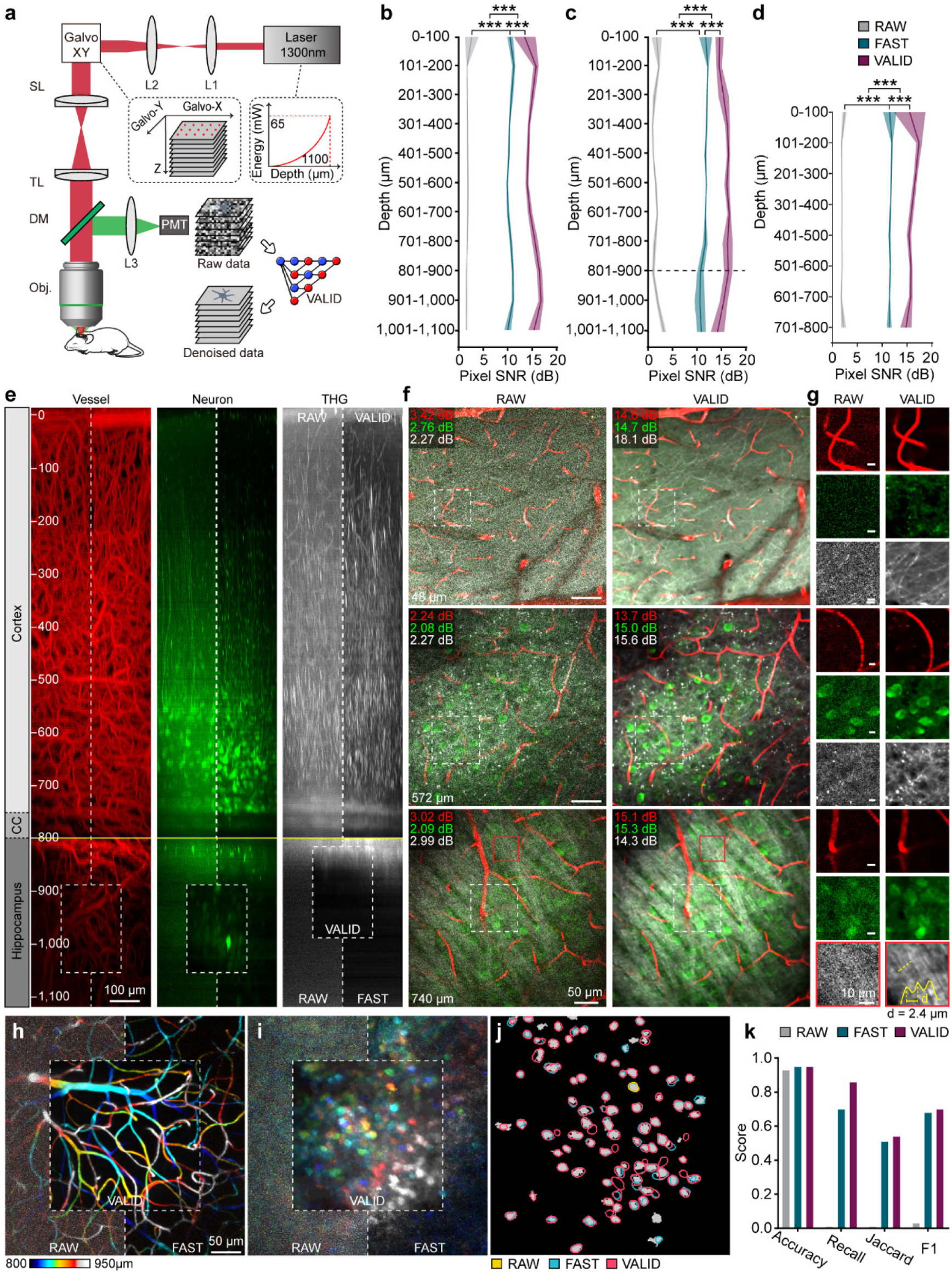
VALID-enhanced three-photon multimodal structural imaging and quantitative evaluation. **a,** Schematic of the three-photon microscope used for intravital imaging. The system employs a Galvo-Galvo (X-Y) scanning configuration and enables synchronized multimodal imaging of vessels (Texas Red), neuronal structures (jGCaMP7s), and third harmonic generation (THG, label-free). **b-d,** SNR plots at different depths for three imaging modalities: blood vessels, neurons, and THG. The SNR values were calculated at various depths, with ground truth (GT) obtained through multi-frame averaging. For each modality, the raw data (RAW), FAST-enhanced results, and VALID-enhanced results are compared. The average SNR values for RAW, FAST, and VALID are as follows: blood vessels (1.47 ± 0.22 dB, 10.37 ± 0.30 dB, 14.97 ± 0.68 dB), neurons (1.30 ± 0.32 dB, 11.33 ± 0.47 dB, 15.49 ± 0.75 dB), and THG (2.03 ± 0.18 dB, 11.65 ± 0.32 dB, 15.96 ± 0.84 dB). Statistical significance is indicated as ***P < 0.001 (unpaired two-tailed t-test). **e,** MIP images along the x-y plane for the three imaging modalities (vessels, neurons, and THG) across depths ranging from 0 to 1,100 μm. The raw data exhibit high noise and low contrast, while VALID-enhanced results show improved structural clarity and continuity. Scale bar, 100 μm. **f,** Axial section views of three representative layers from **e** at cortical depths of 48 μm (superficial layer), 572 μm (intermediate layer), and 740 μm (deep layer). The images demonstrate the fusion of vascular, neuronal, and THG modalities. Scale bar, 50 μm. **g,** Zoomed-in views of ROI from **f**. Most ROIs are indicated with dashed boxes for vascular and neuronal structures, while a specific ROI for THG imaging is marked with a red solid box. For vascular ROIs, VALID results show reduced fragmentation and improved trajectory continuity. For neuronal ROIs, VALID enhances dendritic continuity and somatic differentiation. For THG ROIs, VALID reveals fine neuropil architecture, including intervals of ∼2.4 μm between individual nerve fibers in the corpus callosum. Scale bar, 10 μm. **h,** Depth-encoded MIP images of blood vessels from 900 to 1,050 μm along the z-axis. The depth is color-coded, and VALID results show clearer vessel trajectories compared to RAW and FAST. Scale bar, 50 μm. **i,** Depth-encoded MIP images of neurons from 900 to 1050 μm along the z-axis. The depth is color-coded, and VALID results exhibit enhanced neuronal differentiation compared to RAW and FAST. **j,** Neuronal segmentation results using Cellpose^18^ for RAW, FAST, and VALID images, with GT manually delineated. Segmented neurons are shown in different colors for comparison. **k,** Quantitative evaluation of segmentation performance, including Accuracy, Recall, Jaccard, and F1 scores. VALID achieves the highest values across all metrics.

Despite the high noise level and low contrast of the three-photon data, as the MIP results shown in Fig. 2e, the vessels, neuronal structures, and brain tissue became clear after denoising. The denoising process significantly enhanced structural continuity and interpretability across all modalities. We selected three typical layers with cortical depths of z = 48 μm (superficial layer), 572 μm (intermediate layer), and 740 μm (deep layer), presents an axial section view of the fusion of vascular neurons and THG imaging (Fig. 2f). The zoom-in region of interests (ROIs; Fig. 2g) show that blood vessels demonstrated improved trajectory continuity with reduced fragmentation, forming more coherent vascular networks; the neuronal structures revealed substantial improvement in neuronal signal-to-noise ratio and contrast, particularly evident through enhanced dendritic continuity and superior somatic differentiation that enabled precise demarcation of individual neuronal somas; simultaneously, the THG results (red ROI in Fig. 2g) displaying subtle textures and morphological features in brain tissue, allowing clear visualization of fine neuropil architecture, delineating the intervals between two individual nerve fibers as ∼2.4 μm (yellow profile) in corpus callosum. This comprehensive enhancement collectively validated the denoising algorithm’s capability to resolve critical biological details despite challenging initial imaging conditions.

To further illustrate the superior resolving power for capturing deeper vessels and neurons in the hippocampus, Fig. 2h and 2i show the depth-encoded projection results from 900 to 1050 μm. Undoubtedly, VALID demonstrates superior performance in both modalities compared to RAW and FAST, particularly in terms of neuron differentiation. Moreover, as shown in Fig. 2j, we manually delineated the GT for these neurons in Fig. 2i and then used Cellpose^18^ to segment the raw, FAST, and VALID results, respectively. The results demonstrated that our approach achieved the best performance, which was further supported by quantitative metrics (Fig. 2k). Notably, VALID-denoised data enabled the identification of 104 neurons — a dramatic improvement over the single neuron segmented from the raw data.

### Improving reconstruction efficiency of light-field imaging

Light-field microscopy (LFM) is renowned for its capability of rapid volumetric imaging by simultaneously capturing spatial and angular information^5,40^. As a latest variant, confocal scanning light-field microscopy (csLFM^41^) enables high-speed XYZ-T subcellular volumetric imaging. By integrating line-confocal illumination with a synchronized camera rolling shutter, csLFM achieves sustained 3D biological dynamics recordings while selectively capturing in-focus signals to suppress background fluorescence and minimize photobleaching. However, when observing photosensitive cells with limited excitation photons, csLFM suffers from severe shot noise with degraded image quality, necessitating pre-reconstruction denoising to mitigate crosstalk in densely labeled samples. Our processing workflow begins with channel reorganization of the acquired volume stack (Fig. 3a). The original XYU-T configuration is restructured into XYT-U format to better preserve signal independence within each spatial-angular view. Subsequently, we perform VALID denoising on each XYT volume. The denoised volume stack then undergoes inverse channel reorganization to obtain the pre-reconstructed stack. Finally, a deep learning-based reconstruction^2^ is applied to generate high signal-to-noise ratio, high-resolution 3D time-series data, ensuring enhanced visualization of dynamic biological processes (Supplementary Video 1).

**Figure 3.**
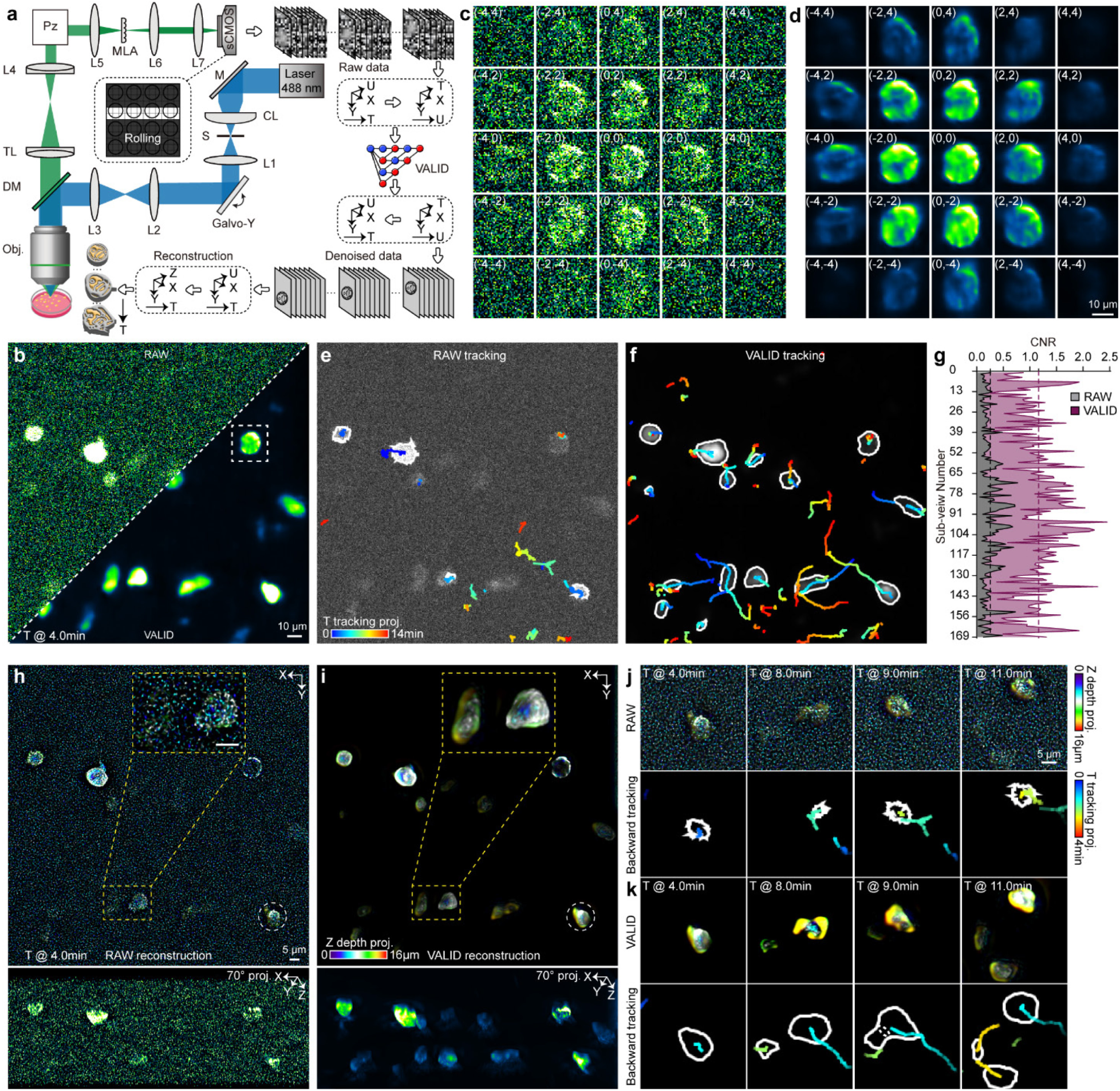
Workflow validation and performance comparison of VALID-enhanced csLFM imaging. **a,** Schematic of the csLFM-VALID processing workflow. The original XYU-T data is reorganized into XYT-U format to preserve signal independence within each spatial-angular view. VALID denoising is then applied to each XYT time-series volume to suppress the shot noise. Subsequently, the denoised stack is restored to its original format through inverse channel reorganization, followed by deep learning-based reconstruction to generate high-SNR 3D time-series data. **b,** Noise reduction comparison at the central spatial-angular image. The raw data (left) exhibits significant noise, whereas the VALID-denoised result (right) shows improved contour clarity. Scale bar, 10 μm. **c,** Raw data from the 5×5 sub-view array. The selected region from the white ROI in **b** shows noticeable noise across multiple spatial-angular views. **d,** VALID-denoised results for the 5×5 sub-view array. Compared to **c**, the VALID-denoised array demonstrates consistent noise suppression and signal preservation across multiple spatial-angular views. Scale bar,10 μm. **e,** Tracking trajectories reconstructed from raw data. The pseudo-color lines represent the trajectories of amoebas over 14 minutes, showing discontinuities and unclear contours. **f,** VALID-denoised tracking trajectories. Compared to **e**, the trajectories exhibit improved continuity and better delineation of amoeba contours. **g,** Quantitative comparison of contrast-to-noise ratio (CNR). The CNR values of raw data (blue line, average 0.256) are significantly lower than those of VALID-denoised data (red line, average 1.166), representing an improvement of approximately 455%. Dashed boxes indicate the locations of three signal ROIs and one background ROI used for CNR calculation. **h,** Reconstruction from raw data with Z-axis depth projection. The yellow ROI highlights a region with blurred subcellular structures, indicating the limitations of raw data reconstruction in revealing fine details. Scale bar, 5 μm. **i,** VALID-denoised reconstruction with Z-axis depth projection. Compared to **h**, the VALID-denoised result reveals clear subcellular features. The inset shows a 70° tilt projection along the x-direction, highlighting significant enhancement in volume reconstruction. **j,** Zoomed-in ROIs and tracking lines at key time points from raw data. The selected frames at 4-, 8-, 9-, and 11-minutes show discontinuous tracking and unclear cellular contours. Scale bar, 5 μm. **k,** Zoomed-in ROIs and tracking lines at key time points after VALID denoising. Compared to **j**, the VALID-denoised results demonstrate improved tracking continuity and enhanced clarity of cellular structures, as indicated by contour lines.

To investigate the enhancement of VALID on the csLFM imaging paradigm, we continuously captured approximately 14 minutes of footage of AX2 axenic strain cells (over-expressing RFP-myosin II) using 13×13 spatial-angular views. These cells are highly photosensitive, requiring ultralow excitation light intensity for long-term volumetric monitoring, which inevitably results in severe shot noise contamination. The comparison between raw data and VALID-denoised results at the central view demonstrates a significant improvement in noise suppression (Fig. 3b). Furthermore, we selected 5×5 spatial-angular views from the rectangular ROI region marked in Fig. 3b. The denoising results across multiple angular views, presented in Fig. 3c and 3d, reveal that VALID achieves excellent performance consistently across all perspectives. To quantify the enhancement, we randomly selected three signal ROIs and one background ROI, calculating the contrast-to-noise ratio (CNR) across all 169 spatial-angular views. Quantitative analysis (Fig. 3g) shows that our method effectively improves the CNR by approximately 455% (0.256 to 1.166, CNR scores), further validating the efficacy of the proposed approach. Fig. 3e and 3f demonstrated the whole 14 minutes cell tracking and projected these courses of events as pseudo-color lines. Notably, the result after denoised by VALID can better delineate the contour of amoebas and achieve more continuous tracking for them, which enables better event extraction.

We further performed 3D reconstruction using both denoised and original data (Fig. 3h and 3i). By applying pseudo-color projection of Z-axis depth (indicated by the yellow ROI regions), it can be clearly observed that the reconstruction results after VALID denoising effectively overcomes the noise bottleneck, unveiling fine subcellular structures that remain obscured in the raw data. To further assess the volumetric enhancement, we performed a 70° tilt projection along the x-axis, demonstrating that VALID significantly enhances the performance across the entire volume. Additionally, Fig. 3j and 3k show the zoom-in ROIs and tracking lines extracted from Fig. 3h and 3i at representative time stamps, respectively. The results clearly demonstrate that after VALID denoising, we can distinctly observe cellular movement, intercellular interactions, and alterations in subcellular structures, features that remain indistinguishable in reconstructions from raw data. These findings collectively prove that our proposed VALID method effectively enhances the conventional csLFM imaging paradigm, enabling long-term high-quality 3D subcellular-resolution imaging with low phototoxicity.

### Breaking through the limitation of speckle noise in optical coherence tomography

Optical Coherence Tomography (OCT^42^) enables non-invasive, high-resolution tissue imaging through low-coherence interferometry. While established in ophthalmology, its dermatological application faces inherent limitations, including the skin’s turbid nature, which amplifies coherent speckle noise caused by multiply-scattered photons. Unlike quantum-limited shot noise (reducible through signal averaging), speckle noise degrades image contrast, obscures microstructural details, and resists conventional denoising due to its structure-dependent characteristics^43^. Current suppression strategies require either computationally intensive algorithms compromising temporal resolution or hardware modifications increasing system complexity^44,45^. VALID introduces self-supervised orthogonal learning and three-dimensional low-frequency Hessian regularization to mitigate speckle artifacts without such trade-offs, addressing a critical barrier to OCT’s advancement in dermatological diagnostics.

To validate the de-speckle capabilities for OCT, we utilized a custom-built spectral-domain OCT (SD-OCT) system to perform volumetric acquisitions of milk and murine skin, where time-series B-scan averaging served as a high SNR reference. In Supplementary Fig. 4, we first prove that VALID performs superior de-speckle performance on homogeneous medium (OCT imaging of milk). Then, we further validate the performance on complex structures (murine skin). Both the raw data, FAST denoised, VALID denoised, and high-SNR Y-Z cross-sectional results at 1050 μm are presented (Fig. 4b). Magnified ROIs reveal the exceptional speckle suppression capability of VALID, with significantly improved tissue clarity compared to other methods. Compared with FAST, VALID not only maintains the uniform background structure but also eliminates speckle artifacts and shows more details. Furthermore, we quantitatively analyzed the spatial frequency characteristics by calculating the power spectrum of ROIs (Fig. 4d) and deriving their optical transfer functions (OTFs; Fig. 4e). The OTF profiles in Fig. 4e demonstrate substantial enhancement across both low- and high-frequency components. Notably, the high-frequency performance of our processed data closely approximates that of the high-SNR reference, confirming the effective preservation of fine structural details. The X-Y cross-section at z = 880 µm in Fig. 4c demonstrates a remarkable isotropic enhancement enabled by VALID, benefiting from orthogonal learning. Through analysis of cross-sectional lines within these ROIs, our approach shows that orthogonal learning simultaneously maintains superior X-axis and Y-axis denoising capabilities while achieving enhanced spatial uniformity, outperforming spatiotemporal sampling-based learning methods (FAST).

**Figure 4.**
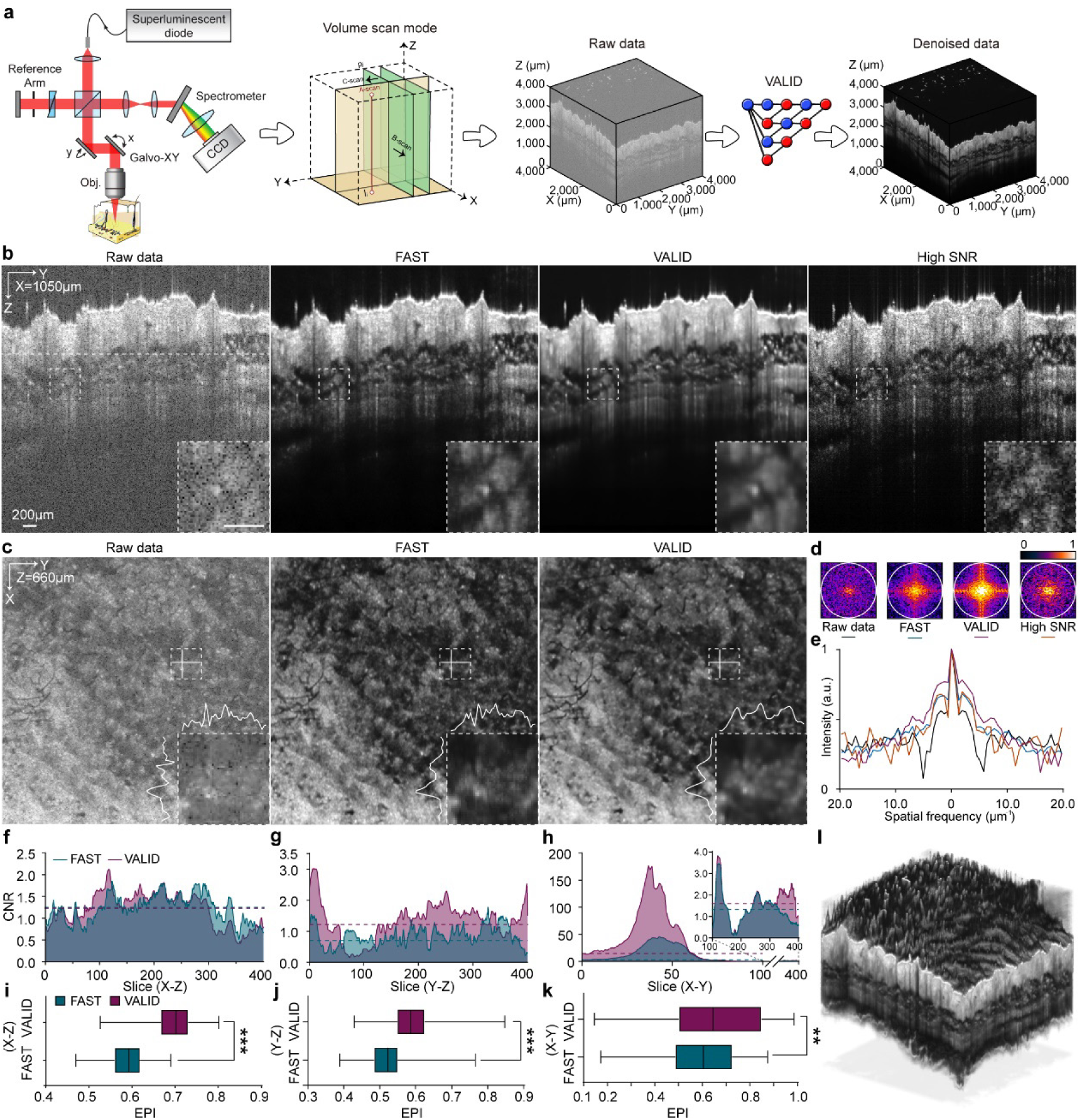
VALID in skin OCT imaging: denoising performance and quantitative validation. **a,** Schematic of the spectral-domain OCT (SD-OCT) imaging system and VALID processing pipeline. The system employs a broadband laser source, beam splitter (BS), reference mirror (M), and spectrometer with a camera to capture interferometric signals. Scanning modes include A-scan (depth-resolved), B-scan (cross-sectional), and C-scan (volumetric). C-scan acquisitions generate 3D raw data volumes, which are subsequently denoised using VALID to suppress speckle noise and enhance tissue clarity. **b,** Comparison of Y-Z cross-sectional images (x = 1,050 μm) across raw data, FAST processing, VALID processing, and a high-SNR reference. VALID achieves superior speckle suppression and tissue clarity, as demonstrated in the magnified ROI. The ROI highlights VALID’s ability to preserve microstructural details compared to other methods. Scale bar, 200 μm. **c,** X-Y cross-sectional images (z = 880 μm) showing isotropic enhancement enabled by VALID. VALID effectively balances denoising along both X and Y axes, improving spatial uniformity compared to raw data. The results demonstrate the benefits of orthogonal learning in achieving consistent noise reduction across planar dimensions. **d,** Power spectrum analysis of the ROI in **b**. Spatial frequency distributions are compared for raw data, FAST, VALID, and the high-SNR reference. VALID closely approximates the power spectrum of the high-SNR reference, particularly in the high-frequency range, indicating effective preservation of fine structural details. **e,** Optical transfer function (OTF) comparison based on the power spectrum in **d**. The dashed line represents the high-SNR reference. VALID demonstrates the highest fidelity to the reference in both low- and high-frequency components, outperforming FAST in preserving high-frequency information critical for microstructural resolution. **f,** CNR analysis in the X-Z plane. VALID achieves comparable performance to FAST in this primary input orientation, demonstrating effective noise suppression without compromising contrast. **g,** CNR analysis in the Y-Z plane. VALID significantly improves CNR by 61.2% compared to raw data, highlighting its ability to enhance structural contrast in orthogonal planes through its self-supervised learning framework. **h,** CNR analysis in the X-Y plane. VALID achieves a remarkable 343.7% improvement in CNR compared to raw data, showcasing its superior performance in isotropic contrast enhancement across planar dimensions. **i,** Edge preservation index (EPI) analysis in the X-Z plane. VALID demonstrates a 17.8% improvement (***, P < 0.001) over existing methods, confirming its ability to retain macroscopic contrast features and microscopic anatomical details. **j,** EPI analysis in the Y-Z plane. VALID achieves a 12.9% enhancement (***, P < 0.001) compared to FAST, further validating its structural fidelity in orthogonal planes. **k,** EPI analysis in the X-Y plane. VALID outperforms FAST with a 7.0% improvement (**, P < 0.01), demonstrating consistent edge preservation across all volumetric dimensions. **l,** 3D rendering of the entire volume after VALID denoising. The reconstruction highlights VALID’s ability to resolve the longstanding trade-off between noise reduction and spatial resolution, achieving a balance that enhances both macroscopic and microscopic tissue features in dermatological OCT imaging.

To provide quantitative validation of VALID’s superiority in contrast preservation and structural detail retention beyond SNR enhancement, we implemented systematic evaluations using two established metrics: contrast-to-noise ratio (CNR) and edge preservation index (EPI). We conducted comprehensive volumetric assessments across three mutually orthogonal planes, sagittal (X-Z), coronal (Y-Z), and axial (X-Y). Comparing contrast performance, an intriguing observation emerges: FAST demonstrates comparable CNR values to VALID in the X-Z plane (its primary input orientation), while VALID achieves significant CNR improvements of 61.2% and 343.7% in Y-Z and X-Y planes respectively, attributable to its orthogonal learning framework that enhances 3D structural consistency (Fig. 4f-4h). The EPI analysis (Fig. 4i-4k) further substantiates VALID’s advantages, with 17.8% (***, P<0.001), 12.9% (***, P<0.001), and 7.0% (**, P<0.01) enhancements over existing methods in X-Z, Y-Z, and X-Y planes correspondingly. These quantitative results systematically demonstrate VALID’s superior capability in preserving both macroscopic contrast features and microscopic anatomical details across full volumetric data. Finally, Fig. 4l shows the rendering result of the whole volume denoised by VALID, demonstrating that VALID addressed the fundamental trade-off between noise reduction and spatial resolution in OCT.

## Discussion

The development of VALID represents a significant advancement in addressing the persistent challenges of volumetric biomedical imaging by establishing a self-supervised orthogonal learning framework that harmonizes structural fidelity, noise suppression, and computational efficiency. Notably, VALID’s self-supervised paradigm overcomes the impracticality of acquiring paired clean-noisy dataset in biomedical imaging, a limitation that has long constrained supervised methods. By leveraging volumetric redundancy through orthogonal planes rather than temporal sequences, the framework maintains compatibility with single-volume acquisitions, a crucial feature for clinical workflows and dynamic process imaging. The significant improvements in deep region neuron detection rates (from 1 to 104 neurons) and depth-independent SNR enhancement (improving ∼10 fold) validate its potential to transform quantitative analysis pipelines, enabling more accurate segmentation, tracking, and biomarker quantification. In addition, VALID’s zero-shot capability enables it to perform high-performance denoising using a single dataset, providing a powerful tool for imaging expensive volumetric electron microscopes (Supplementary Fig. 5).

VALID stands out as a distinctive and powerful denoising technology tailored for high-dimensional biomedical imaging. Unlike conventional denoising methods that primarily depend on temporal redundancy or spatial interpolation, VALID harnesses the inherent three-dimensional structural integrity of volumetric data through its proprietary Tetris sampling protocol and orthogonal cyclic consistency framework. Benefitting from these approaches, VALID has been assigned the perception field covering the entire volume and enables the exploitation of orthogonal cross-planar correlations (X-Y, X-Z, Y-Z) to disentangle anisotropic noise patterns while preserving ultrastructural details, which is particularly critical for imaging modalities operating at quantum noise limits, such as multiphoton microscopy. The incorporation of a CRN further distinguishes VALID from traditional encoder-decoder architectures by prioritizing low-pass filtering in latent space over semantic feature extraction, thereby aligning network design with the low-dimensional nature of denoising tasks. This architectural refinement not only reduces parameter complexity but also enhances interpretability by eliminating redundant hierarchical connections that may propagate high-frequency artifacts.

## Limitations

While VALID establishes new benchmarks, certain limitations warrant consideration. The current implementation employs patch-based processing to manage memory constraints, which introduces computational overhead during volumetric recombination. Future iterations could explore hybrid architectures integrating network operator^46,47^ and quantitative pruning^30,48^ techniques to reduce the requirement of memory for further global contextual modeling. Additionally, as VALID relies on volumetric redundancy, its performance on highly dynamic processes (e.g., voltage signal) remains over-smooth, while this scenario that FAST^31^ is good at. In the future, we will combine the temporal weighted attention mechanism to solve this problem.

## Conclusion

The quantitative and qualitative superiority of VALID across diverse volumetric imaging modalities underscores its versatility in addressing domain-specific noise challenges. In multiphoton microscopy, where nonlinear signal scaling exacerbates depth-dependent noise, VALID achieves unprecedented preservation of subcellular features (e.g., axonal branches < 3 µm) while simultaneously suppressing non-stationary Poisson-Gaussian mixtures, a critical advancement for deep-tissue neuroimaging. For OCT applications, the three-frequency low-frequency Hessian regularization term effectively decouples speckle noise from true tissue textures, addressing the longstanding trade-off between speckle suppression and resolution loss. The framework’s ability to enhance light-field reconstruction efficiency further demonstrates its capacity to synergize with computational high-dimensional imaging paradigms, suggesting broader applicability beyond conventional microscopy.

By achieving zero-shot denoising without sacrificing structural authenticity, VALID lays the foundation for intelligent imaging systems that bridge the gap between acquisition hardware and analytical algorithms. Its success across modalities with fundamentally different noise characteristics (quantum-limited, coherent, and scattering-induced) positions it as a universal enhancer for next-generation volumetric imaging in research and clinical arenas. Finally, based on the above pipelines, we believe our open-source multi-threaded GUI, which incorporates visualized training and inference workflows for maximizing user-friendliness and practicality, will profoundly benefit the biomedical imaging community.

## Methods

### The self-supervised orthogonal learning strategy

We have constructed a self-supervised orthogonal learning strategy for structure-friendly denoising.

Let *y* ∈ ℝ^*X*×*Y*×*Z*^ denote a noisy 3D volume. To generate spatially correlated sub-volumes for self-supervised training, we partition *y* into non-overlapping 2×2×2 blocks.

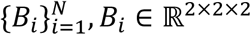

Each block is further divided into four adjacent sub-volumes using orthogonal sub-sampler, named Tetris sampler (as shown in Fig. 1a, denoted as *t*). Then, we can obtain 8 kinds of sub-block as a sampling pool, which is not overlap between central voxels (are orthogonal, denoted as 𝒞).

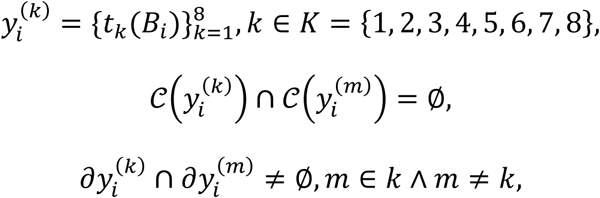

with *∂* denoting voxel boundaries. Next, we randomly selected four sub-blocks for each block *B* and final reorganized as four sub-volumes for training. Formally:

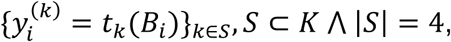

where 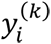 represents the *k* -th sub-volume, and adjacent sub-volumes share overlapping or neighboring voxels in 3D space. By sliding window operation for each *i*, we can obtain the whole volumes *y*_*k*_. This design preserves local structural correlations between sub-volumes, enabling the model to learn denoising from spatial continuity. Meanwhile, the design that makes the centrosomes orthogonal to each other can better eliminate random noise.

For the training framework, we assume the noisy observation *y* follows an additive noise model:

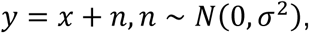

where *x* is the underlying clean volume, and *n* is zero-mean Gaussian noise with variance *σ*^2^. For sub-volumes generated by *G*, we further assume the noise in different sub-volumes *n*_*k*_ is mutually independent and the clean sub-volumes *x*_*k*_ = *g*_*k*_(*x*) retain spatial continuity, i.e., adjacent sub-volumes share consistent structures. Thus, each sub-volume satisfies:

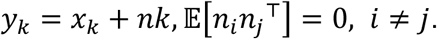

The denoising model *f*_*θ*_ is trained to predict one sub-volume from another, leveraging noise independence. For all pairs (*i*, *j*) where *i* ≠ *j*, we define:

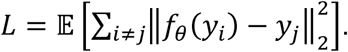

Expanding the expectation:

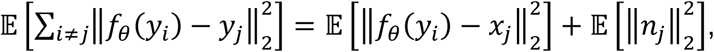

where the cross-term 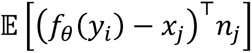 vanishes due to noise independence.

Minimizing *L* forces *f*_*θ*_(*y*_*i*_) to approximate *x*_*j*_, implicitly learning denoising through spatial correlations between *x*_*i*_ and *x*_*j*_. Then the expectation of reconstruction loss is equivalent to supervised learning:

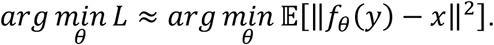

### Network architecture and loss function

We propose a novel cross-scale recursive network architecture to implement VALID, named CRN. As shown in Supplementary Fig.1, when the input is forward into CRN, a conv3d layer with 1×1×1 kernel is employed to extract the features. Then, the feature tensor is down-sampled using AvgPool3d layer for four branches with different scales (↓ 1, ↓ 2, ↓ 4, ↓ 8). For each branch, we design EnhancedConvBlock3d module with adaptive attention (as shown in Supplementary Fig. 1). It is composed of ReflectionPad3d layer, Conv3d layer with 3×3×3 kernel, GroupNorm layer with 4 group number, ELU activation layer, and double repeat them as a EnhancedCovBlock3d block. It is noted that we performed a EnhancedCovBlock3d and then employed the AvgPool3d for each branch. Through cross-scale recursive strategy, the features x^(3,0)^, x^(2,0)^, x^(1,0)^, x^(2,1)^, x^(1,1)^ and x^(1,2)^ are up-sampled using trilinear-based interpolate3d layer. Finally, we concatenated these multi-scale features and using the other conv3d layer with 1×1×1 kernel to format the output.

Our loss function consists of pixel-wise L2-norm loss as the optimization term and our well-designed 3D low-pass Hessian matrix regularization term, which can be formed as following:

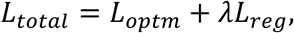

where *L*_*optm*_ presents the optimization term consisting of denoising and identity losses and *L*_*re*_*_g_* notes the regularization term; *λ* is a weighted parameter, we set as 0.0001 here.

The optimization term can be summarized as following:

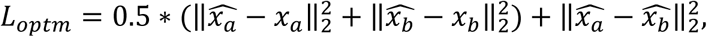

where *x*_*a*_ and *x*_*b*_ are the input sub-volumes; 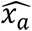 and 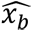 are the output predictions (in Fig. 1a). 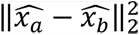 is employed as an identity consistency constraint term.

For regularization term, based-on our previous experience^49^ we first perform three-dimensional wavelet transform to obtain the inherent features of eight different frequencies {H^x^H^y^H^z^, H^x^H^y^L^z^, H^x^L^y^H^z^, H^x^L^y^L^z^, L^x^H^y^H^z^, L^x^H^y^L^z^, L^x^L^y^H^z^ and L^x^L^y^L^z^}. We concatenated the low-frequency features {H^x^L^y^L^z^, L^x^H^y^L^z^, L^x^L^y^H^z^ and L^x^L^y^L^z^}, and then employed three-dimensional Hessian matrix regularization on it to against specific signal-related noise (such as speckle noise). Formally:

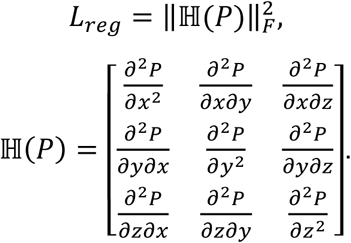

where ℍ denotes the Hessian matrix; P presents the concatenated feature which is composed of two set low-frequency features obtained from 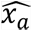 and 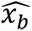 using three-dimensional wavelet transform.

### Training and inference details

VALID was implemented using PyTorch and trained on an NVIDIA RTX A6000 GPU with 48GB memory. The input patch size was fixed at 224×224×224 patch size to accommodate high-resolution 3D data. The complete training dataset (100% of available samples) was utilized without validation set separation. Training was conducted over 100 epochs with a batch size of 1. The Adam optimizer was employed with an initial learning rate of 0.0001 and momentum parameters 0.9 and 0.999. A cosine annealing strategy was adopted for learning rate scheduling, dynamically adjusting the rate throughout the training cycle to enhance convergence stability. For inference, a sliding window approach with 10% overlap along all three axes (x, y, z) was implemented to mitigate boundary artifacts and ensure seamless volumetric predictions. The random seed is 3407. Additionally, we developed a graphical user interface (GUI) for both training and testing pipeline of VALID, positioning it as a novel and effective user-friendly tool applicable for enhancing the conventional volumetric biomedical imaging paradigm. (Supplementary Fig. 6 and Video 2)

### Sample preparation

All experimental procedures were approved by the Animal Welfare and Ethics Committee of the Department of Experimental Animal Science, Fudan University (Ethics Approval Number: 2024-FAET-005).

#### Mouse preparation

All mouse were maintained under controlled environmental conditions (12:12 h light-dark cycle) with *ad libitum* access to food and water.

*For two-photon imaging*, adult transgenic *Thy1-GFP-M* mouse (provided by the Prof. Fang’s laboratory from College of Basic Medical Sciences, Shanghai Jiao Tong University, China) aged 8-10 postnatal weeks were anesthetized using 1.5% isoflurane, and cranially immobilized in a stereotaxic instrument (RWD Life Science, China). Following skull exposure, we used a cranial drill to remove part of the skull and a custom-made coverslip to fit the shape of the cranial window (∼4 mm in diameter) by dental cement. Postoperative analgesia and monitoring are provided. After surgery, mice were housed in a separate cage for 2 weeks of postoperative recovery. For imaging session, animals were re-anesthetized with 3% isoflurane and then fixed onto a custom-made holder.

*For three-photon imaging*, wild-type C57BL6/J mice (8-10 weeks postnatal) received stereotaxic injections of pAAV-hSyn-jGCaMP7s-WPRE (200 nL/site) via glass micropipettes (RWD Life Science, China) into coordinates: AP-2.0 mm, ML-1.76 mm, DV-0.4/-0.8/-1.2/-1.6 mm. After injection, mice were housed in a separate cage for 2-3 weeks of postoperative recovery. When mouse needed to implement cranial window, the operation is the same as two-photon imaging animal preparation. Additionally, to label vessel, we injected 0.1 ml of Texas Red (25 mg/ml, dextran conjugate, 70 kDa, Thermo Fisher Scientific, USA) via tail vein before imaging.

#### Dictyostelium discoideum cell culture

AX2 axenic strain cells were provided by the Professor Cai’s laboratory (National Laboratory of Biomacromolecules, Institure of Biophysics, Chinese Academy of Sciences, China). The AX2 cell line, overexpressing RFP-myosinII, was cultured in HL5 medium (Formedium # HLF2), supplemented with antibiotics, at 22℃. The confocal dish with glass bottom was treated with plasma cleaning, and then coating poly-L-lysine (PLL, 1 μg/ml) in BSS (10mM NaCl, 10mM KCl, 3mM CaCl_2_) for 1h. The cells were washed with BSS 3 times. The movie was captured immediately after transferring the cells onto the PLL surface.

For OCT imaging, wild-type C57BL/6J mice (8-10 weeks postnatal) were sacrificed through anesthetic overdose using a combination of tiletamine-zolazepam (Zoletil®, 150-200 mg/kg) and xylazine (30-40 mg/kg) administered via intraperitoneal (i.p.) injection. Following confirmation of death (via cardiac cessation), the inguinal region underwent depilation using electric clippers followed by depilatory cream application. The subjects were then positioned in a supine orientation on the imaging stage for cutaneous structural acquisition using spectral-domain optical coherence tomography (SD-OCT).

### Imaging system

The experimental setup comprised a multiphoton microscope supporting both two- and three-photon modalities, a custom-built spectral-domain optical coherence tomography system, and a custom-built confocal scanning light-field microscope.

#### Multi-photon microscope

Two- and three-photon *in vivo* volumetric imaging was performed using a commercial multi-photon microscope (DeepVision, MicroLux, China). A specially designed optical system provided a 1.6 mm×1.6 mm field of view (FOV) for both galvo scanning at 0.5 Hz and high-speed resonant scanning at 30 Hz, paired with a 16× water-immersion objective (CFI75 LWD 16X W, Nikon). A near-axial, high-efficiency fluorescence detection path with a collection angle of up to 15 degrees was used to maximize the collection of both ballistic and scattered fluorescence, utilizing high-sensitivity GaAsP PMTs (H10770PA-40, Hamamatsu).

Two-photon imaging was performed using a 920 nm femtosecond pulsed laser (ALCOR 920-2, Spark Lasers) with 100-fs pulse duration and 80-MHz repetition rate. Three-photon imaging employed a 1,300 nm femtosecond pulsed laser (Cronus3P, Light Conversion) with 50-fs pulse duration and 1-MHz repetition rate. To simultaneously acquire third harmonic generation (THG), green fluorescent protein (GFP), and vasculature (Texas Red) signals, emitted photons were spectrally separated into blue (460±30 nm), green (510±30), and red (605±35 nm) channels, respectively. Volumetric imaging was performed over a depth range of 0–1300 µm with a 4 µm inter-plane interval, using a motorized objective stage. Each plane was captured at a 512×512-pixel resolution. Laser power was dynamically adjusted (5–60 mW) to compensate for signal attenuation at increasing depths.

#### Confocal scanning light-field microscope (csLFM)

The inverted csLFM system was employed to capture the 3D dynamics of AX2 axenic strain cells, with detail in ref^41^. A high-NA oil-immersion objective lens (Zeiss Plan-Apochromat ×63/1.4 Oil M27), a high-sensitivity sCOMS camera (Andor Zyla 4.2 PLUS), and a commercial MLA (RPC Photonics MLA-S100-f21) were configurated into the system for 3D imaging with high spatiotemporal resolution. The illumination optical path consisted of multi-channel lasers (Coherent, OBIS 405/488/561/640), a cylindrical lens (Thorlabs, LJ1363L2-A), an optical galvo (Thorlabs, GVS211), and a customized slit with the specific width of 325 μm. During the acquisition of one image, a galvo translates the illumination beam, synchronized with the exposure of rolling shutter, for suppressing the background intensity caused by out-of-focus fluorescence or autofluorescence. After image acquisition, the csLFM images are reconstructed into 3D volumes using deep-learning methods.

#### Optical coherence tomography (OCT)

The study employed a custom-built spectral-domain OCT system. The system integrates a broadband super-luminescent diode (BLM2-D-840-B-10, Superlum, China) with 830 nm central wavelength (±50 nm bandwidth), providing cellular-scale axial resolution essential for detecting epidermal pathological alterations. The spectrometer consisted of a transmission diffraction grating with 1435 lines/mm (Wasatch Photonics, USA), an imaging lens with a focal length of 100 mm, and a line-scan camera (EV71YO1CCL2210-BB3, Teledyne e2v, 2048 pixels, 10 μm pixel width, USA). This configuration achieved 0.051 nm/pixel spectral resolution with 3.4 mm maximum imaging depth in air. The sensitivity roll-off measured 3 dB at 1 mm and 6 dB at 1.6 mm subsurface depth, maintaining sufficient signal-to-noise ratio (SNR) for dermal imaging applications. This system achieves a 5 μm axial resolution and a 10 μm lateral resolution. For each lesion site, we collected one or more three-dimensional volumetric datasets, with the number depending on the size of the lesion. Each OCT volumetric dataset has a FOV of 5 mm by 5 mm and a pixel resolution of 1000 by 1000 with an acquisition time of 10 s.

### Cell segmentation, cell tracking and three-dimensional visualization details

#### Cell segmentation

Maximum intensity projection was applied to the fluorescence time-lapse images to preserve neuronal signals as much as possible, generating single-frame images for subsequent segmentation. The pre-trained Cellpose model^18^ was used for segmentation with the following parameter configurations: cytoplasm segmentation mode was selected, the grayscale channel was used, the nuclear channel was excluded, and the average cell diameter was set to 20 pixels. Ground truth annotations were manually generated using ImageJ’s wand tool by clicking on each cell and adjusting the tolerance to ensure that the selected ROI accurately matched the cell boundaries.

#### Cell tracking

Cell profile was performed using the TraceMate plugin^50^ in ImageJ. A threshold-based segmentation algorithm was applied to distinguish cellular regions from the background. The optimal threshold value was determined automatically by the plugin, leveraging intensity histogram analysis of the input fluorescence or brightfield microscopy images. Morphological operations were subsequently employed to refine segmentation masks and eliminate artifacts. Cell tracking was implemented via the Kalman filter algorithm integrated into the TraceMate plugin. The Kalman filter combined dynamic motion prediction with observed positional data to resolve cell trajectories across sequential time-lapse frames. This approach minimized tracking errors caused by temporary occlusion, overlapping cells, or abrupt motion. Trajectories were validated manually to ensure biological relevance, particularly for mitotic events or complex migration patterns.

#### Three-dimensional visualization

Three-dimensional visualization of segmented cells was achieved using Matlab. Raw 3D image stacks (e.g., z-stack time series) were rendered into volumetric reconstructions, with adjustable parameters for opacity, lighting, and depth-cueing to enhance spatial perception.

#### Depth pseudo-color encoding

It was applied to highlight dynamic cellular behaviors (e.g., migration speed or lineage relationships) using ImageJ’s Temporal Color Code tool. Royal-LUT mapped temporal progression, enabling intuitive visualization of chronological data in composite images.

### Method comparison

We compared the performance of the proposed VALID with four different state-of-the-art comparisons, including SUPPORT^26^, DeepCAD-RT^30^, SRDTrans ^24^, and FAST^31^. For the methods designed for temporal data, we perform the depth axis as time axis, and split the XY-Z image stack into a series of 2D frames to inference the whole volume frame-by-frame. We followed the default framework and network architectures for all methods. The training and inference details are listed in Supplementary Table 1.

### Evaluation metrics

#### Image quality metrics

For the experiments in two-photon microscopy, the reference-based conventional 2D PSNR and SSIM are performed for each frame, which are formulated as follows:

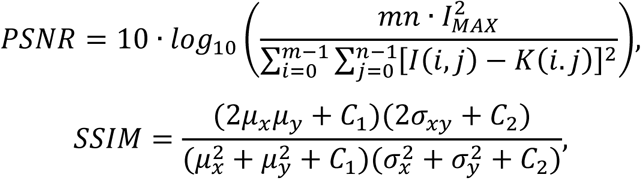

where *I* and *K* present the reference and input image; *I_MAX_* denotes the max pixel value; *m* and *n* are the image width and height. *μ* and *σ* are the means and variances; *C* is constant.

For the experiments in three-photon microscopy, the non-reference-based pixel wised SNR is performed for each frame. Specifically, we perform N imaging operations at each position, then divide each image into d × d sub-regions and calculate the mean value as the result. For each sub-region, we calculate the pixel-level SNR as shown below:

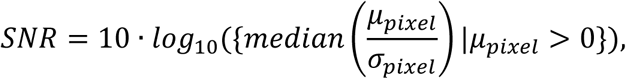

where *μ*_*pixel*_ and *σ*_*pixel*_ are the means and variances of N images at the same layer; we perform median threshold to avoid the extreme conditions at deep region (such as 800 to 1100 μm).

For the experiments in light-field microscopy and OCT, the non-reference-based metric contrast-to-noise ratio (CNR^51^) is performed to demonstrated the contrast and detail preservation. For each sample, we random selected three different signal ROIs and one background ROI to perform three CNR respectively, and then calculate the mean values as the results. The CNR and EPI can be formatted as follows:

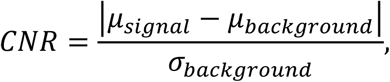

For the experiments in OCT, the non-reference-based metric edge preservation index (EPI^52^) is performed to demonstrated the edge preservation capability. For each sample, we random selected four different ROIs to perform four EPI values, respectively, and then average them as the result. EPI essentially represents the Pearson correlation coefficient of the gradient map, with its value range being [-1, 1]. The closer the value is to 1, the better the edge preservation is and can be formatted as:

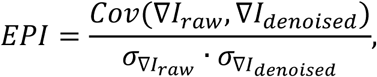

where *I*_*ra*_*_w_* and *I*_*denoised*_ are the original noisy frame and denoised frame; ∇ denotes the Sobel operation; *Cov*(·) presents the covariance; *σ* is the standard deviation.

#### Segmentation metrics

We compared the segmentation results of the raw images and the denoised images with manual ground truth (GT) labels to evaluate the improvement in segmentation brought by the denoising methods. The segmentation results were quantitatively assessed using five metrics: Jaccard, Accuracy, Recall, and F1 score, defined as follows:

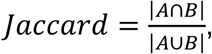

where *A* is the set of predicted pixels, and *B* is the set of ground truth pixels.

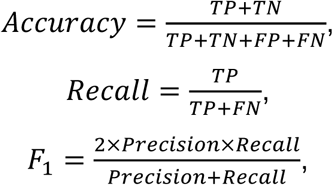

where *TP* is true positives, *TN* is true negatives, *FP* is false positives, and *FN* is false negatives.

## Acknowledgements

This work was supported in part by the National Key R&D Program of China (2022YFF0708700), Shanghai Basic Research Special Zone Program (22TQ020), Natural Science Foundation of Shanghai (22ZR1404300), Shanghai Science and Technology Innovation Action Plan (22S31905500), Medical Engineering Fund of Fudan University (yg2021-032, yg2022-2), Young Scientists Fund of Beijing Natural Science Foundation (4254114), National Postdoctoral Program for Innovative Talent (BX20230174), Postdoctoral Science Foundation of China (2023M741963), Shuimu Tsinghua Scholar Program (2023SM065). The computations in this research were performed using the CFFF platform of Fudan University.

## Data availability

Data are available from the corresponding author upon reasonable request.

## Code availability

The underlying code and GUI software for this study are available at https://github.com/FDU-donglab/VALID.

## References

1. Xu, C., Nedergaard, M., Fowell, D. J., Friedl, P. & Ji, N. Multiphoton fluorescence microscopy for in vivo imaging. Cell 187, 4458–4487 (2024).

2. Lu, Z. et al. Virtual-scanning light-field microscopy for robust snapshot high-resolution volumetric imaging. Nat. Methods 20, 735–746 (2023).

3. Zhao, Z. et al. Two-photon synthetic aperture microscopy for minimally invasive fast 3D imaging of native subcellular behaviors in deep tissue. Cell 186, 2475–2491.e22 (2023).

4. Choe, K. et al. Intravital three-photon microscopy allows visualization over the entire depth of mouse lymph nodes. Nat. Immunol. 23, 330–340 (2022).

5. Wu, J. et al. Iterative tomography with digital adaptive optics permits hour-long intravital observation of 3D subcellular dynamics at millisecond scale. Cell 184, 3318–3332.e17 (2021).

6. Bouma, B. E. et al. Optical coherence tomography. Nat. Rev. Methods Primer 2, 1– 20 (2022).

7. Schubert, M. C. et al. Deep intravital brain tumor imaging enabled by tailored three-photon microscopy and analysis. Nat. Commun. 15, 7383 (2024).

8. Michalska, J. M. et al. Imaging brain tissue architecture across millimeter to nanometer scales. Nat. Biotechnol. 42, 1051–1064 (2024).

9. Scheele, C. L. G. J. et al. Multiphoton intravital microscopy of rodents. Nat. Rev. Methods Primer 2, 1–26 (2022).

10. Icha, J., Weber, M., Waters, J. C. & Norden, C. Phototoxicity in live fluorescence microscopy, and how to avoid it. BioEssays 39, 1700003 (2017).

11. Laissue, P. P., Alghamdi, R. A., Tomancak, P., Reynaud, E. G. & Shroff, H. Assessing phototoxicity in live fluorescence imaging. Nat. Methods 14, 657–661 (2017).

12. Scherf, N. & Huisken, J. The smart and gentle microscope. Nat. Biotechnol. 33, 815–818 (2015).

13. Neifeld, M. A. Information, resolution, and space–bandwidth product. Opt. Lett. 23, 1477–1479 (1998).

14. Wu, J., Ji, N. & Tsia, K. K. Speed scaling in multiphoton fluorescence microscopy. Nat. Photonics 15, 800–812 (2021).

15. Corbetta, E. & Bocklitz, T. Machine Learning-Based Estimation of Experimental Artifacts and Image Quality in Fluorescence Microscopy. Adv. Intell. Syst. 7, 2400491 (2025).

16. Lin, R., Guo, Y., Jiang, W. & Wang, Y. Advanced technologies for the study of neuronal cross-organ regulation: a narrative review. Adv. Technol. Neurosci. 1, 166 (2024).

17. Lefebvre, A. E. Y. T. et al. Nellie: automated organelle segmentation, tracking and hierarchical feature extraction in 2D/3D live-cell microscopy. Nat. Methods 1–13 (2025) doi:10.1038/s41592-025-02612-7.

18. Stringer, C., Wang, T., Michaelos, M. & Pachitariu, M. Cellpose: a generalist algorithm for cellular segmentation. Nat. Methods 18, 100–106 (2021).

19. Chaudhary, S., Moon, S. & Lu, H. Fast, efficient, and accurate neuro-imaging denoising via supervised deep learning. Nat. Commun. 13, 5165 (2022).

20. Weigert, M. et al. Content-aware image restoration: pushing the limits of fluorescence microscopy. Nat. Methods 15, 1090–1097 (2018).

21. Guo, M. et al. Deep learning-based aberration compensation improves contrast and resolution in fluorescence microscopy. Nat. Commun. 16, 313 (2025).

22. Qu, L. et al. Self-inspired learning for denoising live-cell super-resolution microscopy. Nat. Methods 21, 1895–1908 (2024).

23. Qiao, C. et al. Zero-shot learning enables instant denoising and super-resolution in optical fluorescence microscopy. Nat. Commun. 15, 4180 (2024).

24. Li, X. et al. Spatial redundancy transformer for self-supervised fluorescence image denoising. Nat. Comput. Sci. 3, 1067–1080 (2023).

25. Zhang, G. et al. Bio-friendly long-term subcellular dynamic recording by self-supervised image enhancement microscopy. Nat. Methods 20, 1957–1970 (2023).

26. Eom, M. et al. Statistically unbiased prediction enables accurate denoising of voltage imaging data. Nat. Methods 20, 1581–1592 (2023).

27. Ning, K. et al. Deep self-learning enables fast, high-fidelity isotropic resolution restoration for volumetric fluorescence microscopy. Light Sci. Appl. 12, 204 (2023).

28. Lecoq, J. et al. Removing independent noise in systems neuroscience data using DeepInterpolation. Nat. Methods 18, 1401–1408 (2021).

29. Li, X. et al. Reinforcing neuron extraction and spike inference in calcium imaging using deep self-supervised denoising. Nat. Methods 18, 1395–1400 (2021).

30. Li, X. et al. Real-time denoising enables high-sensitivity fluorescence time-lapse imaging beyond the shot-noise limit. Nat. Biotechnol. 41, 282–292 (2023).

31. Dong, B., et al. Real-time self-supervised denoising for high-speed fluorescence neural imaging. Preprint at 10.21203/rs.3.rs-6101322/v1 (2025).

32. Falk, T. et al. U-Net: deep learning for cell counting, detection, and morphometry. Nat. Methods 16, 67–70 (2019).

33. Wang, Z., Xie, Y. & Ji, S. Global voxel transformer networks for augmented microscopy. Nat. Mach. Intell. 3, 161–171 (2021).

34. Jin, L. et al. Deep learning enables structured illumination microscopy with low light levels and enhanced speed. Nat. Commun. 11, 1934 (2020).

35. Olaf Ronneberger, Philipp Fischer, & Thomas Brox. U-Net: Convolutional Networks for Biomedical Image Segmentation. in Medical Image Computing and Computer-Assisted Intervention – MICCAI 2015 vol. 9351 (Springer, Cham, 2015).

36. Helmchen, F. & Denk, W. Deep tissue two-photon microscopy. Nat. Methods 2, 932–940 (2005).

37. Thornton, M. A. et al. Long-term in vivo three-photon imaging reveals region-specific differences in healthy and regenerative oligodendrogenesis. Nat. Neurosci. 27, 846–861 (2024).

38. Horton, N. G. et al. In vivo three-photon microscopy of subcortical structures within an intact mouse brain. Nat. Photonics 7, 205–209 (2013).

39. Ouzounov, D. G. et al. In vivo three-photon imaging of activity of GCaMP6-labeled neurons deep in intact mouse brain. Nat. Methods 14, 388–390 (2017).

40. Prevedel, R. et al. Simultaneous whole-animal 3D imaging of neuronal activity using light-field microscopy. Nat. Methods 11, 727–730 (2014).

41. Lu, Z. et al. Long-term intravital subcellular imaging with confocal scanning light-field microscopy. Nat. Biotechnol. 1–12 (2024) doi:10.1038/s41587-024-02249-5.

42. de Boer, J. F. Spectral/Fourier Domain Optical Coherence Tomography. in Optical Coherence Tomography: Technology and Applications (eds. Drexler, W. & Fujimoto, J. G.) 165–193 (Springer International Publishing, Cham, 2015). doi:10.1007/978-3-319-06419-2_6.

43. Liba, O. et al. Speckle-modulating optical coherence tomography in living mice and humans. Nat. Commun. 8, 15845 (2017).

44. Lee, W. et al. Deep learning-based image enhancement in optical coherence tomography by exploiting interference fringe. Commun. Biol. 6, 1–12 (2023).

45. Mazlin, V. et al. Real-time non-contact cellular imaging and angiography of human cornea and limbus with common-path full-field/SD OCT. Nat. Commun. 11, 1868 (2020).

46. Niu, W., Guan, J., Wang, Y., Agrawal, G. & Ren, B. DNNFusion: accelerating deep neural networks execution with advanced operator fusion. in Proceedings of the 42nd ACM SIGPLAN International Conference on Programming Language Design and Implementation 883–898 (Association for Computing Machinery, New York, NY, USA, 2021). doi:10.1145/3453483.3454083.

47. Cai, X., Wang, Y. & Zhang, L. Optimus: An Operator Fusion Framework for Deep Neural Networks. ACM Trans Embed Comput Syst 22, 1:1–1:26 (2022).

48. Liang, T., Glossner, J., Wang, L., Shi, S. & Zhang, X. Pruning and quantization for deep neural network acceleration: A survey. Neurocomputing 461, 370–403 (2021).

## References

49. Gu, Y., Guan, Y., Yu, Z. & Dong, B. SegCoFusion: An Integrative Multimodal Volumetric Segmentation Cooperating With Fusion Pipeline to Enhance Lesion Awareness. IEEE J. Biomed. Health Inform. 27, 5860–5871 (2023).

50. Tinevez, J.-Y. et al. TrackMate: An open and extensible platform for single-particle tracking. Methods 115, 80–90 (2017).

51. Fang, L. et al. Fast Acquisition and Reconstruction of Optical Coherence Tomography Images via Sparse Representation. IEEE Trans. Med. Imaging 32, 2034–2049 (2013).

52. Geng, M. et al. Triplet Cross-Fusion Learning for Unpaired Image Denoising in Optical Coherence Tomography. IEEE Trans. Med. Imaging 41, 3357–3372 (2022).

