## Supplementary Information for "Enhancing Biomedical Optical Volumetric Imaging via Self-Supervised Orthogonal Learning"

**Supplementary Figure 1:** Network architecture of CRN.

**Supplementary Figure 2:** Evaluating the performance of structure restoration.

**Supplementary Figure 3:** Evaluating the performance of out-of-domain data.

**Supplementary Figure 4:** Evaluating the de-speckle capabilities of OCT using milk.

**Supplementary Figure 5:** The denoising performance of VALID on vEM.

**Supplementary Figure 6:** Python-based cross-platform VALID graphical user interface overview.

**Supplementary Table 1:** List of denoising methods compared in this study.

**Supplementary Table 2:** List of external datasets.

**Supplementary Video 1:** The denoising performance of VALID on photosensitive subcellular recordings captured by csLFM.

**Supplementary Video 2:** The demonstration of VALID’s graphical user interface.

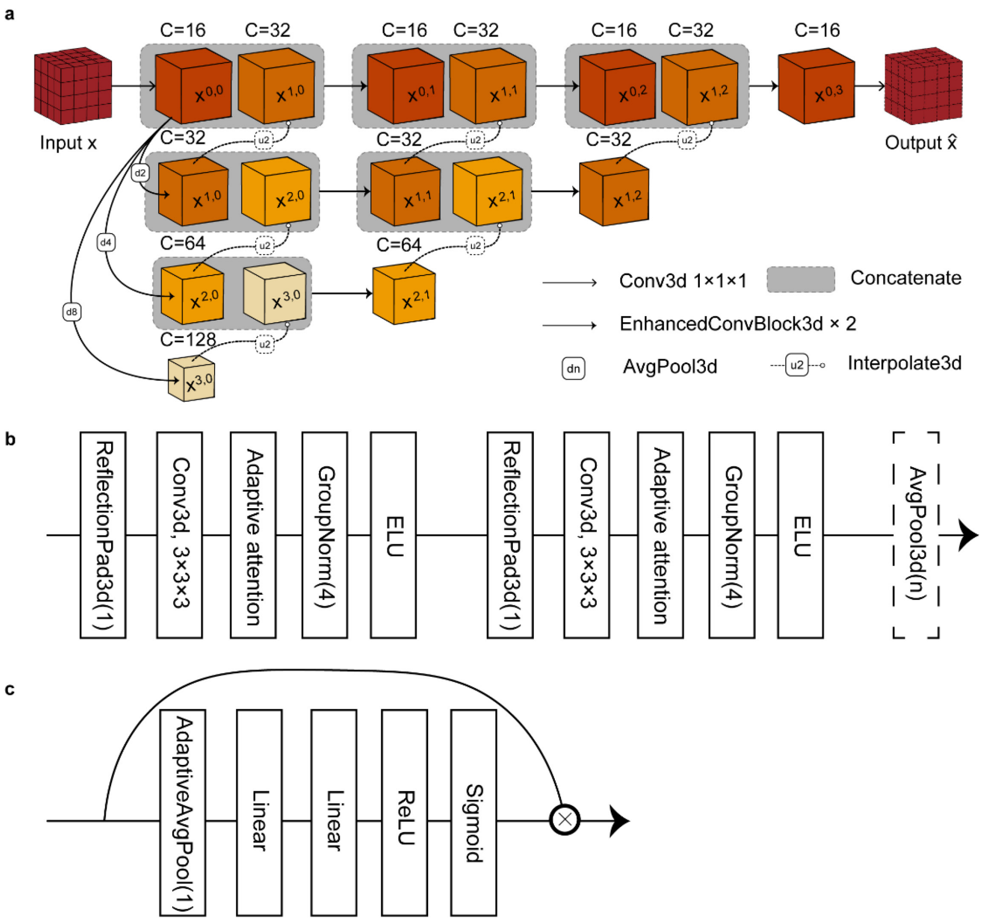


**Supplementary Figure 1: Network architecture of CRN. a**, The overall structure of the network. The input is processed through four different feature scales using EnhancedConvBlock3d for convolution. Features from these scales are concatenated to produce the final output. **b**, The structure of the EnhancedConvBlock3d. This block is executed twice, with each iteration consisting of ReflectionPad, Conv3d, Adaptive Attention, GroupNorm, and ELU. The block is followed by n iterations of AvgPool3d. **c**, The structure of Adaptive Attention. This module implements a channel-wise attention mechanism. Specifically, the input is first globally pooled using AdaptiveAvgPool, then passed through two fully connected (Linear) layers with ReLU activation after the first layer and Sigmoid activation after the second. The resulting attention weights are multiplied element-wise with the original input to produce the output.

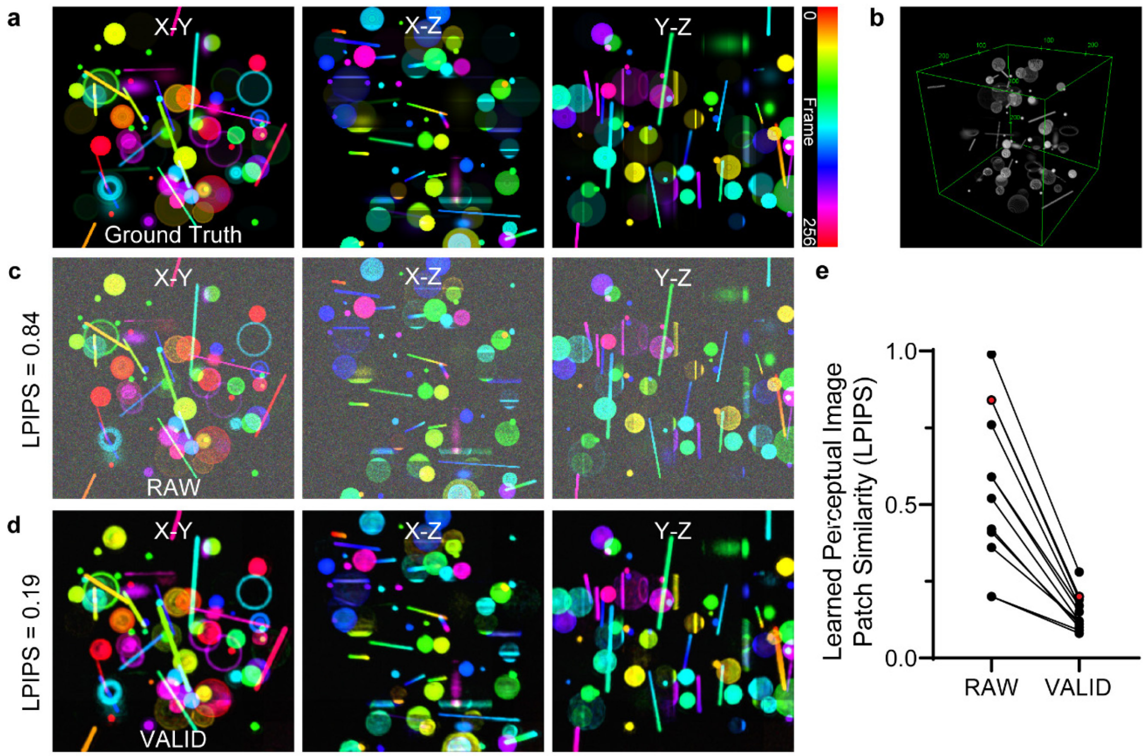


**Supplementary Figure 2: Evaluating the performance of structure restoration. a,** Synthetic volume's ground truth of size 256×256×256 were generated, containing randomly placed solid and hollow spheres, ellipsoids, and rods. The figure shows the max intensity projections of the volumes, where different colors represent varying depths along the axis. **b,** Render result of a. **c,** Noise (lambda=100; sigma=35) was added to simulate data. Learned perceptual image patch similarity (LPIPS) evaluated the maintaining capability of structure. Lower LPIPS value is better. **b,** Results of denoising the noisy data with VALID. Structures are effectively recovered from the noisy input, showing significant improvement after denoising. The figure highlights how VALID reconstructs clear structural details even under low-SNR conditions. **c,** Ground truth of synthetic volume and render result **d,** LPIPS improvement curves for data with different initial levels after denoising with VALID. The plot demonstrates that VALID is robust across a wide range of input LPIPS values, consistently improving the signal quality and achieving effective noise suppression. The red point denotes the sample demonstrated in **a** and **b**.

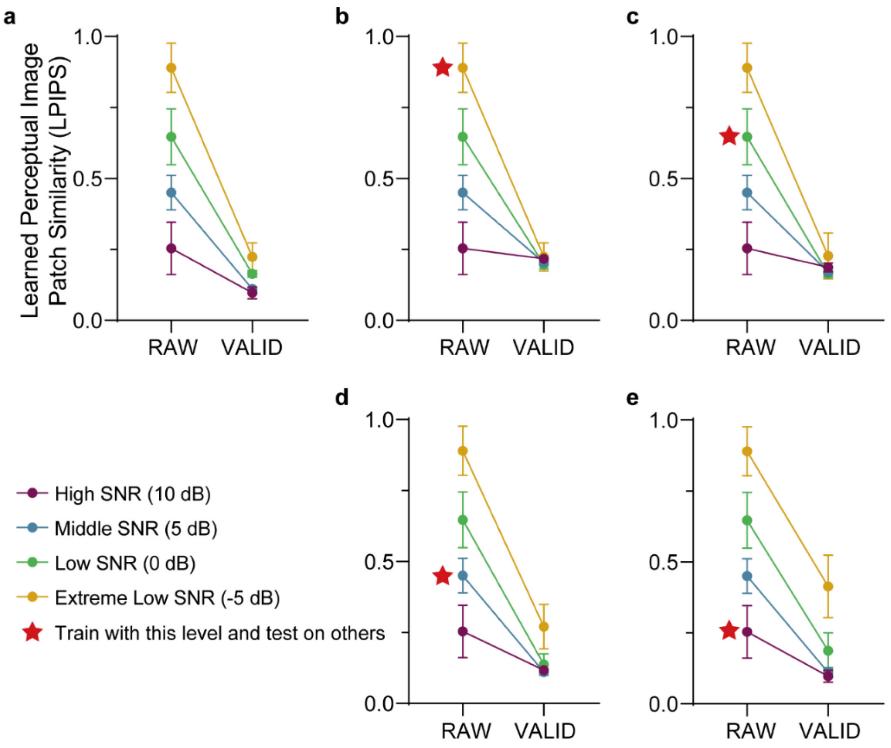


**Supplementary Figure 3: Evaluating the performance of out-of-domain data.** This figure demonstrates the generalizability of the VALID model across varying SNR levels. LPIPS results are presented as mean ± SD. **a,** T The VALID model was trained and tested on datasets with the same SNR levels, showing consistent denoising performance across a wide range of SNRs. **b,** The model was trained on data with a mean SNR of -5 dB and tested on datasets with different SNR levels, achieving robust performance across conditions. **c,** The model trained on 0 dB mean SNR data retained strong denoising capability when applied to datasets with varying SNRs. **d,** When trained on 5 dB mean SNR data, the model demonstrated effective generalization and noise suppression across other SNR levels. **e,** Training on 10 dB mean SNR data resulted in slightly reduced performance for extremely low SNR datasets, but the model remained versatile overall.

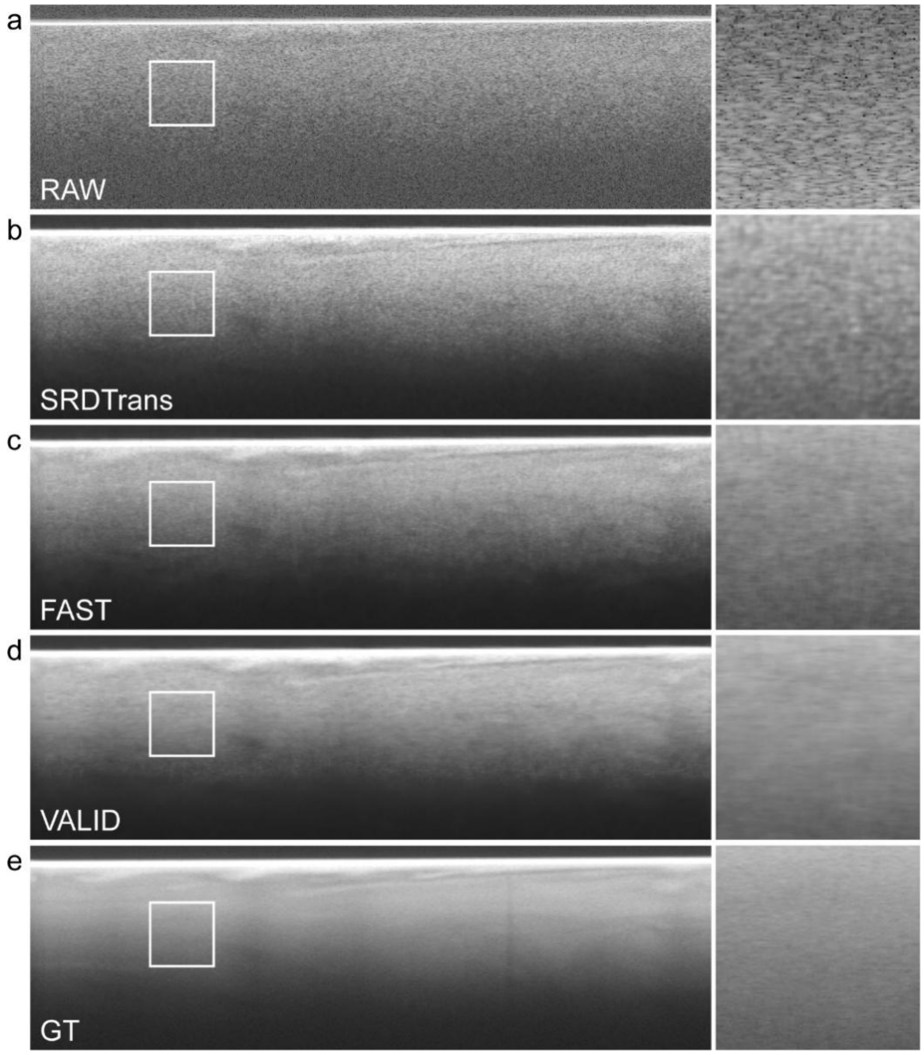


**Supplementary Figure 4: Benchmarking the de-speckle capabilities of OCT.** To evaluate the de-speckle performance of various methods, we used optical coherence tomography (OCT) imaging of diluted milk, which does not contain inherent structural features. The results demonstrate a progressive reduction in speckle noise across different methods. **a,** The original OCT image, showing prominent speckle noise distributed uniformly across the field. **b,** The de-speckled image processed by SRDTrans. Speckle noise is reduced to some extent, but residual noise patterns are still noticeable. c, The de-speckled image processed by FAST. Compared to SRDTrans, FAST achieves further speckle reduction, but some noise artifacts remain visible. **d,** The de-speckled image processed by VALID. VALID shows the most effective speckle suppression, resulting in a smoother and cleaner image compared to other methods. **e,** The ground truth image, representing the ideal speckle-free result. From a to d, the speckle noise exhibits a clear decreasing trend, with VALID achieving the most significant noise reduction. These results highlight the potential of VALID for robust de-speckling in OCT imaging, even in datasets without structural features.

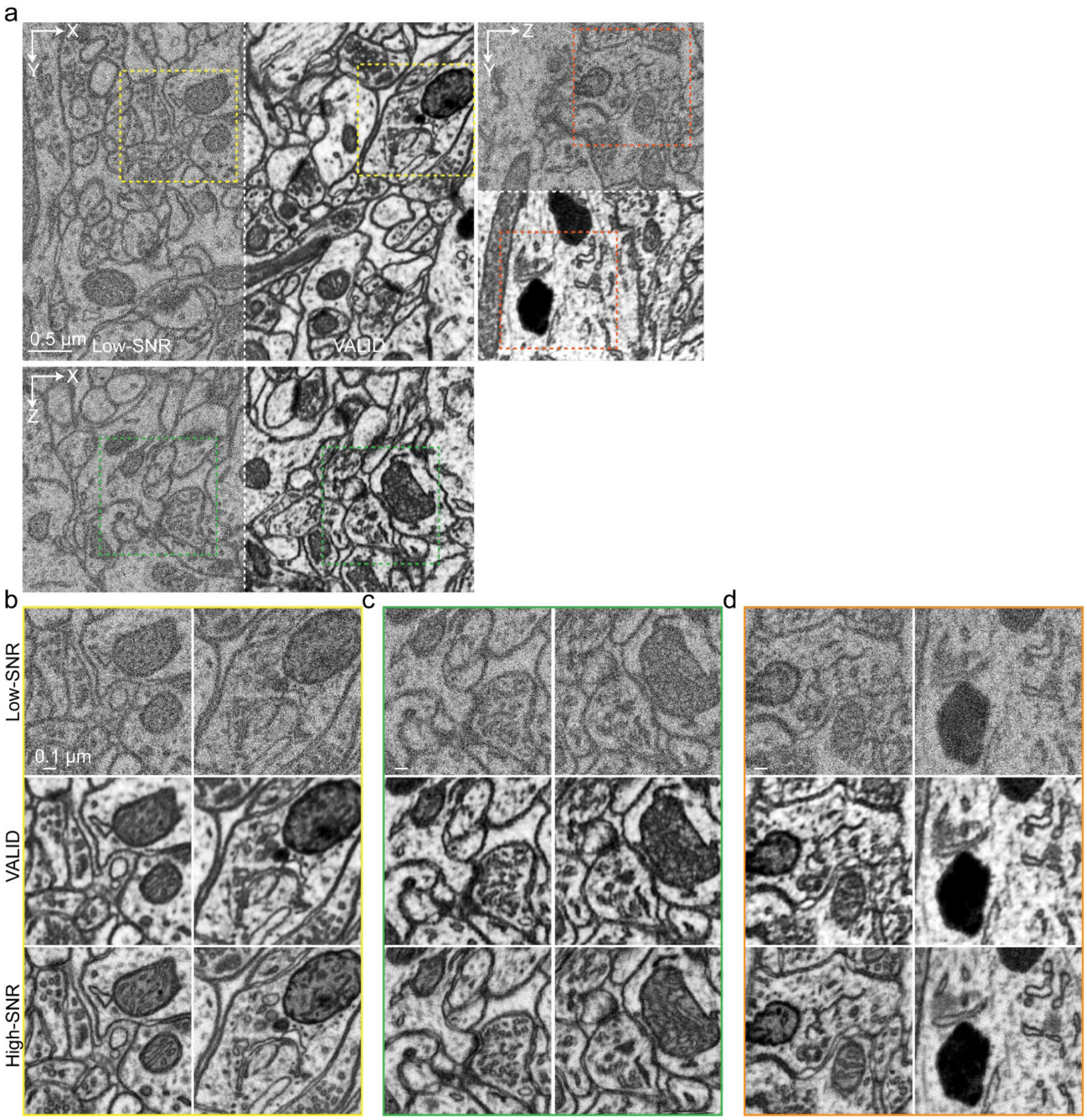


**Supplementary Figure 5: The denoising performance of VALID on vEM.** This figure illustrates the effectiveness of the VALID model in denoising volumetric electron microscopy (vEM) data of a mouse brain, highlighting its ability to enhance image quality while preserving structural details. **a,** Comparison between raw low-SNR vEM data (Supplementary Table 2) and the corresponding VALID-denoised data. The denoised images exhibit significantly improved clarity and reduced noise, enabling better visualization of fine structures within the mouse brain. Scale bar, 0.5 μm. **b,** Zoom-in views of selected regions of interest (ROIs) in the X-Y plane. These magnified areas demonstrate the VALID model's capability to recover intricate cellular and subcellular features that are otherwise obscured by noise in the raw data. Scale bar, 0.1 μm. **c,** Zoom-in views of ROIs in the X-Z plane. The VALID-denoised data reveal improved depth resolution and structural continuity across slices, underscoring the model's utility for analyzing volumetric datasets. Scale bar, 0.1 μm. d, Zoom-in views of ROIs in the Y-Z plane. The results further confirm the VALID model's ability to enhance image quality across different orientations, ensuring accurate representation of 3D biological structures. Scale bar, 0.1 μm.

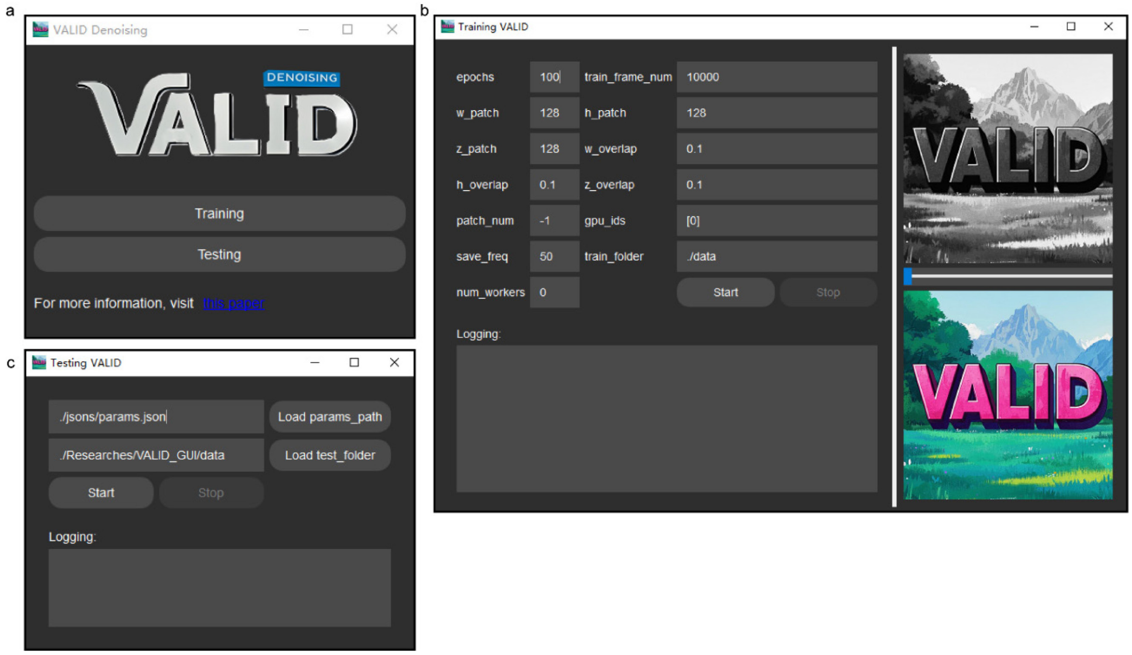


**Supplementary Figure 6: Python-based Cross-platform VALID Graphical User Interface Overview. a,** Main window of VALID's GUI consists of training and testing widgets. **b,** Training pipeline of VALID. **c,** Testing pipeline of VALID. This GUI is compatible with cross-platform usage, including Windows, Linux, and Mac OS.

**Supplementary Table 1:** List of denoising methods compared in this study.

| **Method** | **Repository URL** | **Framework** | **Reference** |
|---|---|---|---|
| SUPPORT | https://github.com/NICALab/SUPPORT | PyTorch | Eom, M. et al. Statistically unbiased prediction enables accurate denoising of voltage imaging data. Nat Methods 20, 1581–1592 (2023). |
| DeepCAD-RT | https://github.com/cabooster/DeepCAD-RT | PyTorch | Li, X. et al. Real-time denoising enables high-sensitivity fluorescence time-lapse imaging beyond the shot-noise limit. Nat Biotechnol 41, 282–292 (2023). |
| SRDTrans | https://github.com/cabooster/SRDTrans | PyTorch | Li, X. et al. Spatial redundancy transformer for self-supervised fluorescence image denoising. Nat Comput Sci 3, 1067–1080 (2023). |
| FAST | https://github.com/FDU-donglab/FAST | PyTorch | Wang, Y. et al. Real-time self-supervised denoising for high-speed fluorescence neural imaging, 16 March 2025, PREPRINT (Version 1) available at Research Square [https://doi.org/10.21203/rs.3.rs-6101322/v1] |

**Supplementary Table 2:** List of external datasets.

| **Method** | **Sample** | **Repository URL** | **Reference** |
|---|---|---|---|
| vEM | Mouse Brain | https://www.epfl.ch/labs/cvlab/data/data-em/ | Lucchi, A. et al. Learning for Structured Prediction Using Approximate Subgradient Descent with Working Sets, 2013 IEEE Conference on Computer Vision and Pattern Recognition, Portland, OR, USA, 1987-1994 (2013) |
